# SosA inhibits cell division in *Staphylococcus aureus* in response to DNA damage

**DOI:** 10.1101/364299

**Authors:** Martin S. Bojer, Katarzyna Wacnik, Peter Kjelgaard, Clement Gallay, Amy L. Bottomley, Marianne T. Cohn, Gunnar Lindahl, Dorte Frees, Jan-Willem Veening, Simon J. Foster, Hanne Ingmer

## Abstract

Inhibition of cell division is critical for cell viability under DNA damaging conditions. In bacterial cells, DNA damage induces the SOS response, a process that inhibits cell division while repairs are being made. In coccoid bacteria, such as the human pathogen *Staphylococcus aureus*, the process remains poorly understood. Here we have characterized an SOS-induced cell-division inhibitor, SosA, in *S. aureus*. We find that in contrast to the wildtype, *sosA* mutant cells continue division under DNA damaging conditions with decreased viability as a consequence. Conversely, overproduction of SosA leads to cell division inhibition and reduced growth. The SosA protein is localized in the bacterial membrane and mutation of an extracellular amino acid, conserved between homologs of other staphylococcal species, abolished the inhibitory activity as did truncation of the C-terminal 30 amino acids. In contrast, C-terminal truncation of 10 amino acids lead to SosA accumulation and a strong cell division inhibitory activity. A similar phenotype was observed upon expression of wildtype SosA in a mutant lacking the membrane protease, CtpA. Thus, the extracellular C-terminus of SosA is required both for cell-division inhibition and for turnover of the protein. Functional studies showed that SosA is likely to interact with one or more divisome components and, without interfering with early cell-division events, halts cell division at a point where septum formation is initiated yet being unable to progress to septum closure. Our findings provide important insights into cell-division regulation in staphylococci that may foster development of new classes of antibiotics targeting this essential process.

**Importance:** *Staphylococcus aureus* is a serious human pathogen and a model organism for cell-division studies in spherical bacteria. We show that SosA is the DNA-damage-inducible cell-division inhibitor in *S. aureus* that upon expression causes cell swelling and cessation of the cell cycle at a characteristic stage post septum initiation but prior to division plate completion. SosA appears to function via an extracellular activity and is likely to do so by interfering with the essential membrane-associated division proteins, while at the same time being negatively regulated by the membrane protease CtpA. This report represents the first description of the process behind cell-division inhibition in coccoid bacteria. As several pathogens are included in this category, uncovering the molecular details of SosA activity and control can lead to identification of new targets for development of valuable anti-bacterial drugs.

## Introduction

Bacteria multiply by coordinated and essential DNA replication and cell-division events, two important biological processes that are valuable targets for antimicrobial therapy (1-4). In the event of DNA damage, the SOS response is activated and ensures that cell division is delayed until the DNA is repaired. The SOS regulon is controlled by the conserved LexA repressor, which is inactivated in response to RecA, a sensor of DNA damage at stalled replication forks, binding to single-stranded DNA (5-7). The SOS response has mostly been studied in *Escherichia coli*, a gram-negative rod-shaped bacterium where the LexA-regulated gene, *sulA*, encodes a cell-division inhibitor. This inhibitor suppresses division by binding to FtsZ, which acts as a scaffold for the assembly of cell-division components at the division site (4, 8-13). In rod-shaped bacteria, inhibition of cell division leads to filamentation as a consequence of lateral peptidoglycan synthesis. This phenotype is also observed in *E. coli* mutants lacking the Lon protease of which SulA is a substrate, further substantiating the role of SulA in cell-division inhibition and filamentation (14, 15).

SulA is poorly conserved in species outside Enterobacteriaceae. The a-proteobacterium *Caulobacter crescentus* encodes an SOS-induced cell-division inhibitor SidA that does not show homology to SulA. In contrast to cytosolic SulA, SidA is a membrane-anchored, small protein that does not interact with FtsZ but rather with later cell-division proteins (16). In this species, yet another, possibly redundant, cell-division inhibitor DidA is implicated in survival during DNA damage, while being independent on the SOS response (17). Hence, it appears that bacteria choose fundamentally different ways to orchestrate regulated cell-division inhibition in response to DNA damage. For gram-positive, spherical cells such as *Staphylococcus aureus*, however, little is known of how SOS induction is coupled to cell division nor have specific posttranslational mechanisms for the negative regulation of cell-division inhibition been identified.

*S. aureus* is a serious gram-positive human pathogen, notorious for being implicated in a wide range of infections and for being able to acquire resistance towards important antibiotic classes. It originally received its name from the grape-like clusters of coccoid cells that result from the unique cell-division process that occurs in three consecutive orthogonal planes (18-20). In this organism, we and others have previously identified *lexA* and noted an open reading frame (designated *sosA*) that is divergently transcribed from *lexA* and controlled by the LexA repressor and the SOS response (21-24). The location of *sosA* adjacent to *lexA* indicated that it might encode a cell-division inhibitor, as similar gene synteny has been observed for SOS-induced, cell-division inhibitors encoded by gram-positive, rod-shaped bacteria, namely *Bacillus subtilis* (25), *Bacillus megaterium* (26), *Listeria monocytogenes* (27, 28), *Mycobacterium tuberculosis* (29), and *Corynebacterium glutamicum* (30) (Fig. 1A). The cell-division inhibitors of these organisms display no similarity to SulA, and within the group they generally show little homology with the exceptions of YneA from *L. monocytogenes* and ChiZ from *M. tuberculosis* (29) that resemble *B. subtilis* YneA. Commonly, however, they have a single membrane spanning segment, a predicted extracellular C-terminus and, except for DivS from *C. glutamicum*, a LysM domain, which is a ubiquitous peptidoglycan binding motif (31, 32), suggesting that membrane and/or cell-wall localization is a common feature characterizing gram-positive cell-division inhibitors.

**Fig 1.**
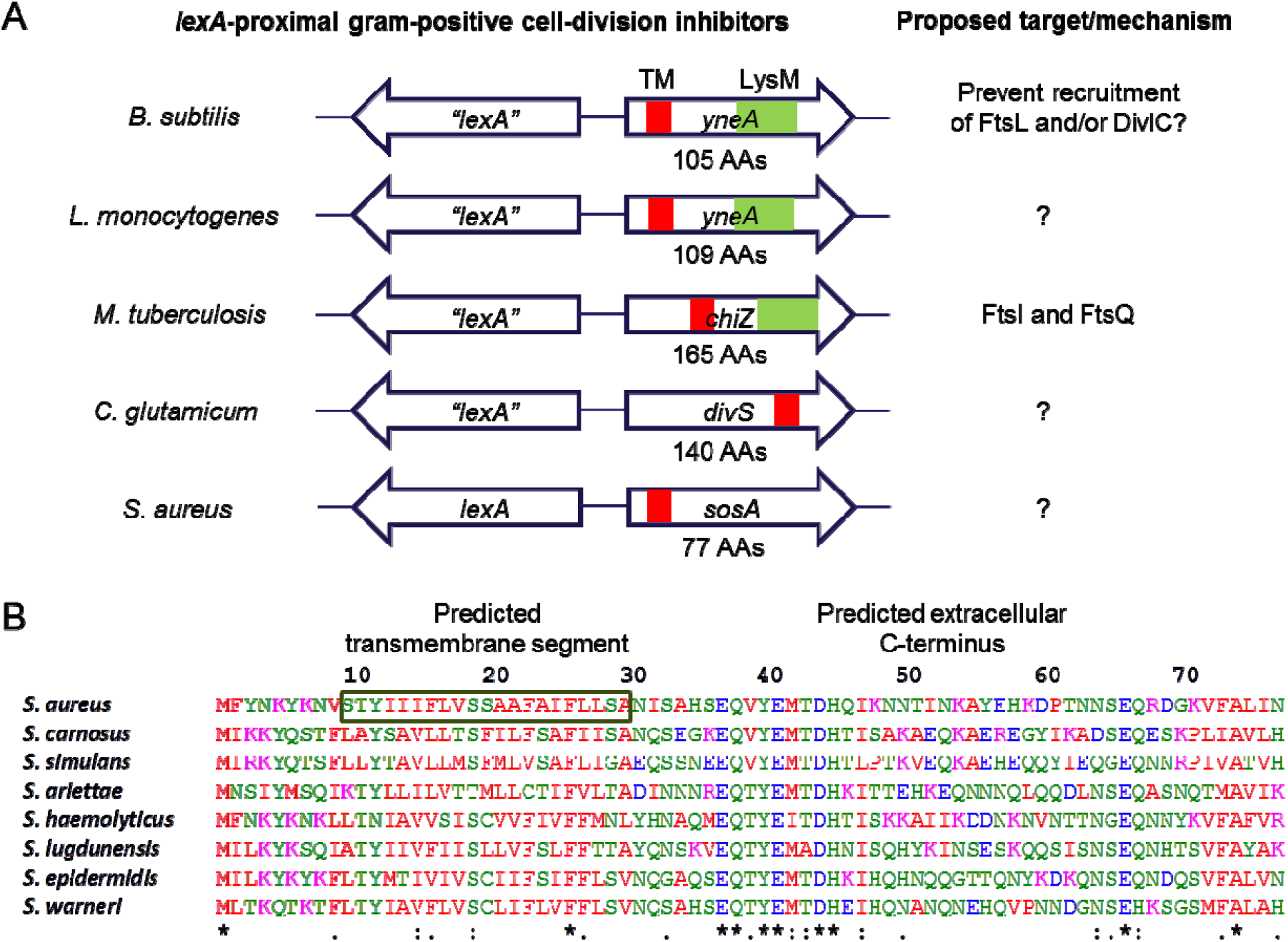
Gram-positive SOS-controlled cell-division inhibitors. (A) Schematic representation of genes encoding characterized gram-positive SOS-regulated cell-division inhibitors (not drawn to scale) including the uncharacterized *sosA* from *S. aureus*. Despite considerable sequence divergence, these genes are commonly chromosomally co-localized with *lexA* homologous genes. The cell-division inhibitors carry a single transmembrane domain (TM), and several proteins have an additional LysM domain. (B) Alignment (CLUSTAL O[1.2.4]) of SosA sequences deduced from open reading frames next to *lexA* in *S. aureus* strain 8325-4 (YP_499864) and seven *Staphylococcus* species: *S. carnosus* (CAL27889), *S. simulans* (AMG96201), *S. arlettae* (EJY94737), *S. haemolyticus* (YP_253482), *S. lugdunensis* (YP_003471776), *S. epidermidis* (YP_188489), and *S. warneri*(EEQ79882). The proteins are 77 amino acids long and are characterized by a predicted transmembrane segment at AAs 10-30 (forS. *aureus* SosA) and a predicted extracellular C-terminal (TOPCONS [37]) with considerable sequence conservation at the membrane-proximal portion (“*” indicates fully conserved residues, “:” indicates conservation of residues with highly similar properties).

The fundamental processes of staphylococcal cell division have been studied spatiotemporally using super-resolution microscopy techniques (19, 33), and recent efforts combining genetic approaches have unveiled the molecular mechanism in unprecedented detail (34, 35). Here, we demonstrate that *sosA* encodes the SOS-inducible cell-division inhibitor in *S. aureus* and document its impact on cell division following treatment with DNA damaging agents. Moreover, we identify a possible mechanism for the proteolytic control of endogenous cell-division inhibition. Thus, we further our insight into these basic biological phenomena, which could lead to the development of new antimicrobials targeting the cell-division machinery in *S. aureus*.

## Results

### Conservation of *sosA* in staphylococci

In *S. aureus*, the open reading frame located adjacent to *lexA* (divergently transcribed) was named *sosA* and hypothesized to encode an inhibitor of cell division (Fig. 1A) (23). The 77-amino-acid-long product has homology and 25-60% amino-acid identity to proteins encoded by genes occupying the same chromosomal location in a range of staphylococci (Fig. 1B and Fig. SI), while showing no clear homology to the known gram-positive cell-division inhibitors. Of note, the C-terminal part of SosA lacks a LysM domain, which is present in the cell-division inhibitors YneA and ChiZ (36, 37). The N-terminal half of SosA includes a putative transmembrane (TM) domain, and the C-terminal part is predicted to be located extracellularly (TOPCONS server [38]). Among the different staphylococcal species, a highly conserved sequence is located just C-terminally of the predicted TM domain and, although less striking, some conservation also seems to exist in the extreme C-terminus (Fig. 1B).

### Genotoxic stress inhibits cell division in an SosA-dependent manner

To analyze whether SosA serves as an SOS-induced cell-division inhibitor, we exposed 8325-4, the common laboratory *S. aureus* strain, and JE2, a derivative of the clinically relevant USA300 lineage, to a lethal concentration of the DNA damaging agent Mitomycin C (MMC) and observed induction of SosA expression in both strains by western blot analysis (Fig. S2A). In contrast, no SosA was observed in deletion mutant derivatives but was expressed in a complemented strain. (Fig. S2A). Upon exposure to the same lethal dose of MMC for a 2 hour period, wildtype cells showed 10-100-fold greater survival than the mutant cells lacking SosA, whereas the optical densities of both cell type cultures were comparable (Fig. 2A). Flow cytometry measurements of cell size distributions revealed that treatment with MMC led to a three- and six-fold increase in forward light scatter (FSC-A) values in 8325-4 and JE2, respectively, while the cell size distribution of the mutants lacking *sosA* remained essentially constant (Fig. 2B). The constant cell size but increasing optical densities of the *sosA* mutant populations implies that during DNA damage these cells, in contrast to wildtype cells, continue to divide. Indeed, this striking phenotype was confirmed by time-lapse microscopy, where wildtype cells increased in size over time, occasionally growing so large that lysis occurred, while the size of the mutant cells lacking *sosA* stayed the same and cell division continued (Fig. 2C and Movie SI). The phenotypes of the 8325-4 *sosA* mutant were abrogated in a strain complemented with a chromosomal copy of *sosA* expressed from its native promoter (Fig. S2B and C). Thus, SosA seems to prevent or delay cell division; maintain viability of *S. aureus* under DNA damaging conditions, and as a consequence of continued peptidoglycan synthesis cause cell size to increase.

**Fig 2.**
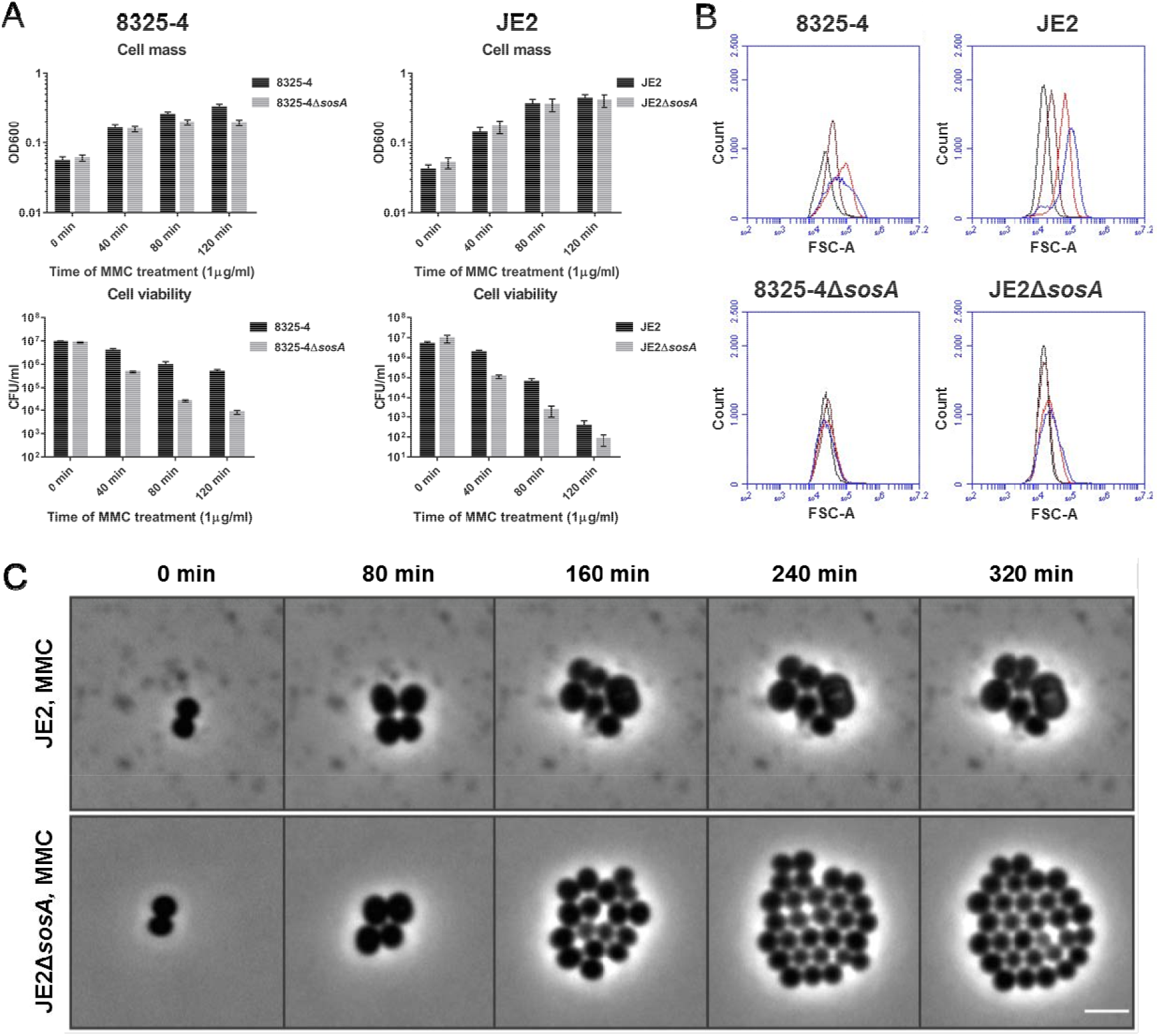
SosA supports survival of *S. aureus* subjected to lethal DNA damage and is involved in bacterial swelling. (A) Culture optical density at 600 nm and cell viability of *S. aureus* strains 83254 and JE2 in comparison with their respective Δ*sosA* mutants upon challenge with a lethal dose of mitomycin C (MMC, 1 μg/ml) for two hours. Error bars represent the standard deviation from three biological replicates. (B) Cell size of 8325-4 and JE2 wildtype and Δ*sosA* mutants exposed to mitomycin C estimated by flow cytometry (FSC-A). Cells were grown exponentially prior to MMC addition at an OD_600_ of 0.05. Samples were taken after 0 (black), 40 (brown), 80 (red), and 120 (blue) minutes of incubation with MMC. (C) Effect of MMC treatment (0.04 μg/ml_) on cell shape and cell number of JE2 wildtype and JE2ΔsosA as visualized by time-lapse phase-contrast microscopy. Scale bar represents 2 μm.

### SosA overexpression inhibits cell division

If SosA is indeed a cell-division inhibitor in *S. aureus*, it can be predicted that overexpression of the protein should be sufficient to inhibit growth even under conditions with no DNA damage. Hence, we expressed *sosA* episomally from an anhydrotetracycline (AHT)-inducible promoter and found that while the control vector did not affect cell viability, the plating efficiency of cells carrying the *sosA* overexpression plasmid was markedly compromised under inducing conditions (Fig. 3A). When grown in liquid culture and compared to the control, the SosA overexpressing cells showed a 2-3-fold increase in forward light scatter values, reflecting that the size of the *sosA*-expressing cells increased (Fig. 3B). Microscopy observations over time confirmed the slight increase in cell size and the reduction in the overall number of cells upon SosA overproduction (Fig. 3C and Movie S2). Interestingly, contrary to what was observed under DNA-damaging conditions (Fig. 2B), we found that ectopic induction of SosA caused only a transient increase in cell size (compare time points 45 and 90 min to 135 min in Fig. 3B), indicating the existence of a negative regulatory mechanism.

**Fig 3.**
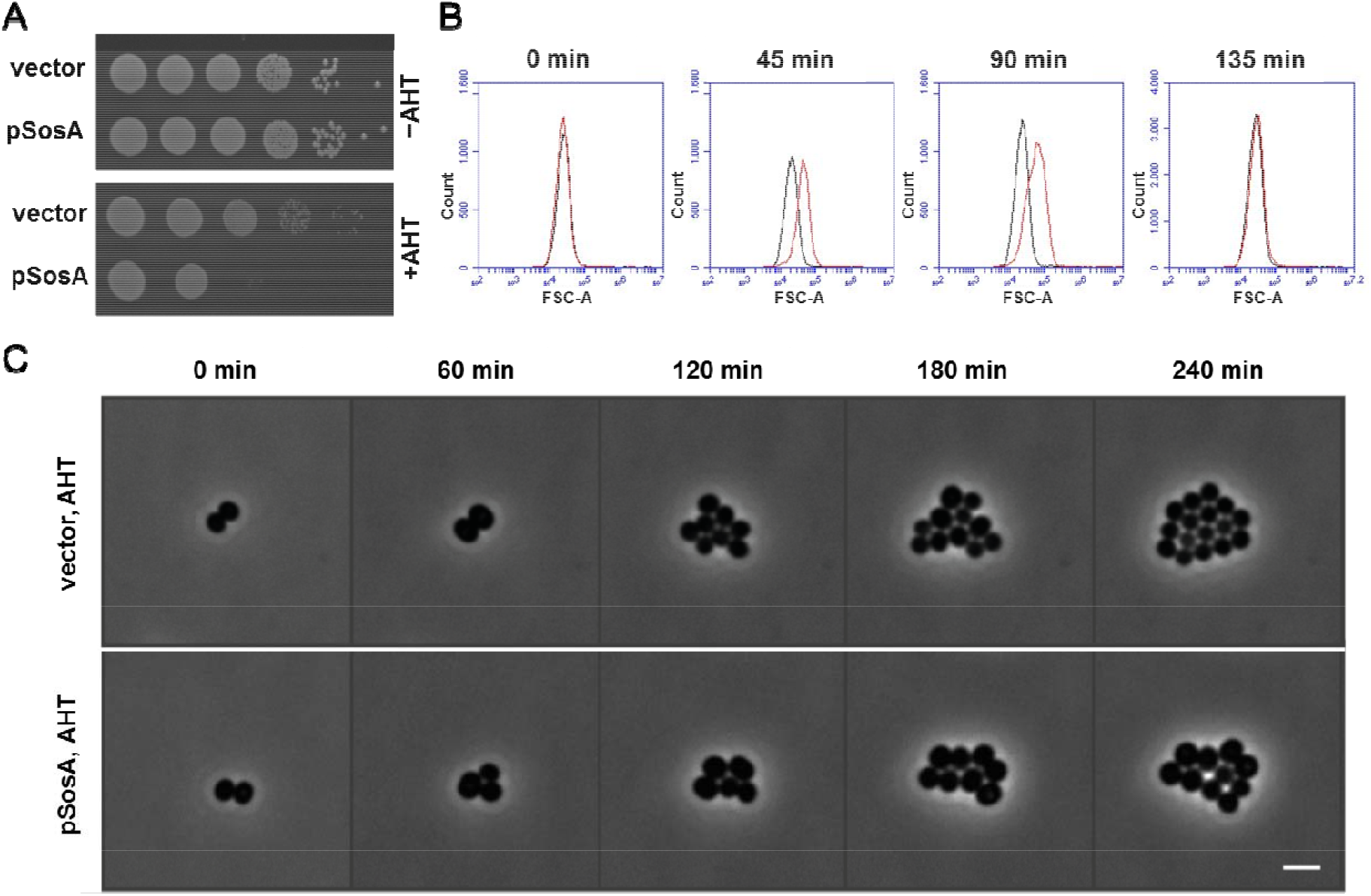
Expression of *sosA* alone interferes with *S. aureus* growth. (A) The effect of a controlled expression of *sosA* on the ability to form colonies was assessed in *S. aureus* RN4220. Strains carrying either the vector control (vector) or *sosA* under an anhydrotetracycline (AHT)-inducible promoter (pSosA) were grown exponentially to an OD_600_ of 0.5, serially 10-fold diluted, and plated on TSA plates in the presence or absence of TSA+300 ng/ml of AHT. The plates were incubated overnight at 37°C and imaged. (B) Evaluation of cell size distribution by flow cytometry (FSC-A) of *S. aureus* RN4220 containing the control vector (black) or pSosA (red). Cells were grown exponentially prior to induction with 100 ng/ml of AHT. At the indicated time points, the cells were collected and analyzed by flow cytometry. (C) Visualization of cell size increase and reduction of cell number of JE2/pRAB12–/ocZ (control) and JE2/pRAB12-sos>4 in the presence of 200 ng/mL of AHT by time-lapse phase-contrast microscopy at 37^ĝ^C. Scale bar represents 2 μm.

### The C-terminal part of SosA is functionally essential and possibly required for autoregulation

The C-terminal part of SosA contains segments that appear somewhat conserved in staphylococci (Fig. 1B), and *in silico* it is predicted to be extracellular. First, we confirmed its extracellular location by constructing PhoA-LacZ reporter fusions (39), whereby a fusion of the reporter chimera to the C-terminus of SosA conferred a phosphatase-positive, ß-galatosidase-negative phenotype in *E. coli*, indicative of translocation across the membrane (Fig. 4). Next, aiming at a functional characterization of its role in SosA activity, we constructed a series of C-terminally truncated variants by the successive elimination of ten amino acids (AAs), resulting in four variants of SosA lacking between 10 and 40 of the C-terminal AAs (Fig. 5A). Strikingly, we observed that truncation of the 10 C-terminal AAs (and to some extent also removal of 20 C-terminal AAs) strongly compromised plating efficiency even at low inducer concentrations (Fig. 5B). Time-lapse microscopy revealed that cells producing SosAd10 (SosA missing the last 10 AAs) had a dramatic phenotype, with prominent cell swelling and a reduction of the cell-cycle rate (Fig. 5C and Movie S3). The phenotypes from the plating assay correlated with the cell size measurements by flow cytometry (Fig. S2D). These results indicate that cells expressing the 10 AA truncated protein are stalled in a non-dividing state leading to abnormal cell size, whereas those expressing full-length SosA are only transiently halted or delayed in cell division (Fig. S2D and Movie S3). In contrast, deletion of 30 or 40 AAs from the C-terminal part eliminated the ability of SosA to inhibit cell division, as cells expressing SosAd30 or SosAd40 gave comparable plating efficiency as those carrying the vector (Fig. 5B) and caused no effect on cell size upon induction (Fig. S2D). This prompted us to investigate the relevance of the highly conserved residues located in the membrane-proximal part of the SosA C-terminal part by performing alanine substitutions within the protein. To this end, we used the highly inhibitory SosAd10 variant as a scaffold and found that while point mutations at AA positions 37 or 38 (E and Q) had no effect, point mutations at positions 40 or 41 (Y and E) partially inactivated and the mutations at positions 44 and 45 (D and H) completely abolished the cell-division inhibitory activity of the protein (Fig. 6A). The mutation at AA position 44 (D to A) was subsequently shown also inactivate the inhibitory activity of the full-length protein (Fig. 6B). Importantly, the inactive variants SosAd40 and SosAd10(44A) are still likely to be localized in the membrane, as are SosA and SosAd10, when evaluated by the PhoA-LacZ translocation assay (Fig. 4). We conclude that conserved residues in the membrane-proximal segment of the extracellular C-terminal part of SosA are essential for inhibiting cell division, while truncation at the extreme C-terminus of the protein augments activity.

**Fig 4.**
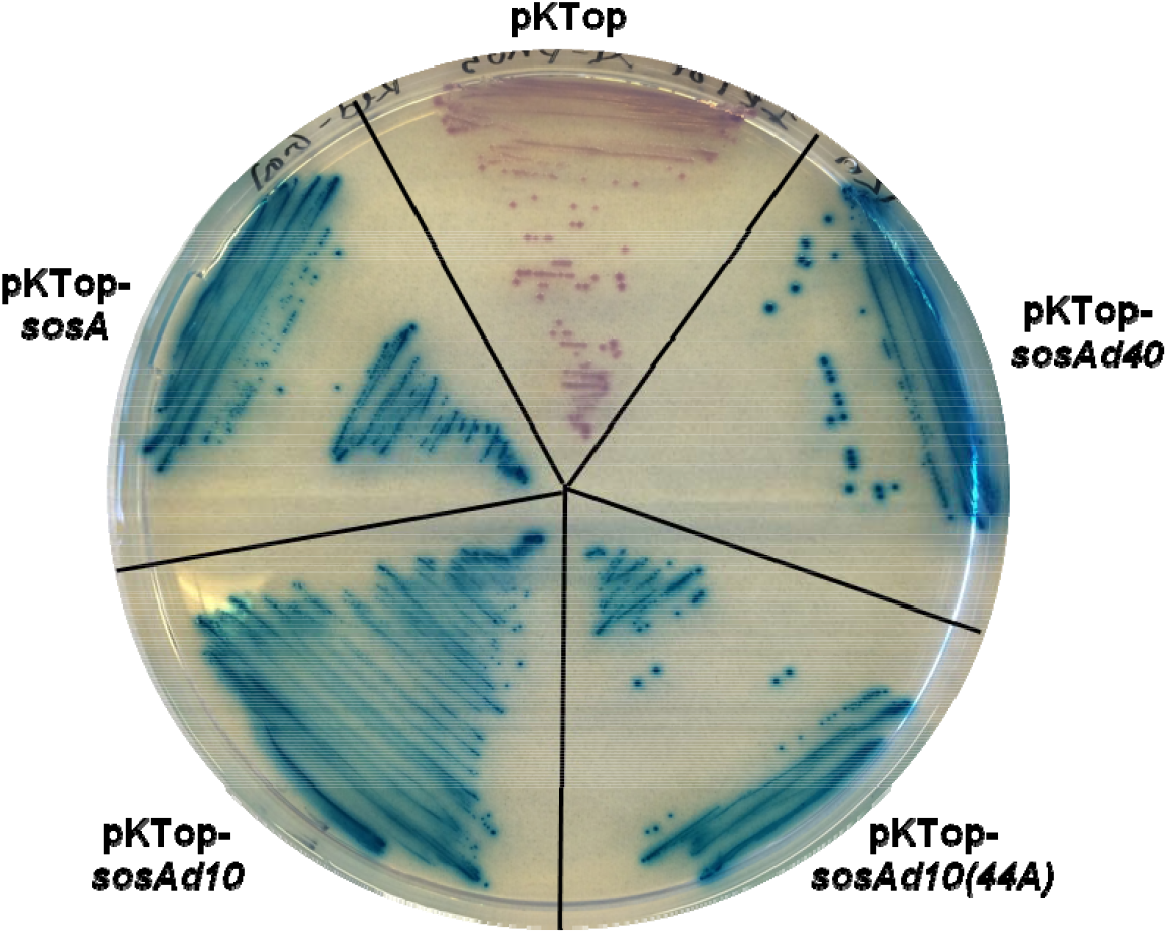
Evaluation of membrane localization of SosA and truncated variants SosAd10, SosAd10(44A), and SosAd40 by in frame fusion at the C-terminus to PhoA-LacZ in pKTop (the truncated variants are explained in Fig. 5 and Fig. 6). *E. coli* IM08B cells carrying the constructs were streaked on a dual indicator plate containing LB agar plus 50 μg/ml of kanamycin, 1 mM IPTG, 5-bromo-4-chloro-3-indolyl phosphate disodium salt (80 μg/ml) and 6-Chloro-3-indolyl-ß-D-galactopyranoside (100 μg/ml). Cytoplasmic localization of the PhoA-LacZ chimera is indicated by red/rose color development whereas translocation across the membrane is indicated by blue color development.

**Fig 5.**
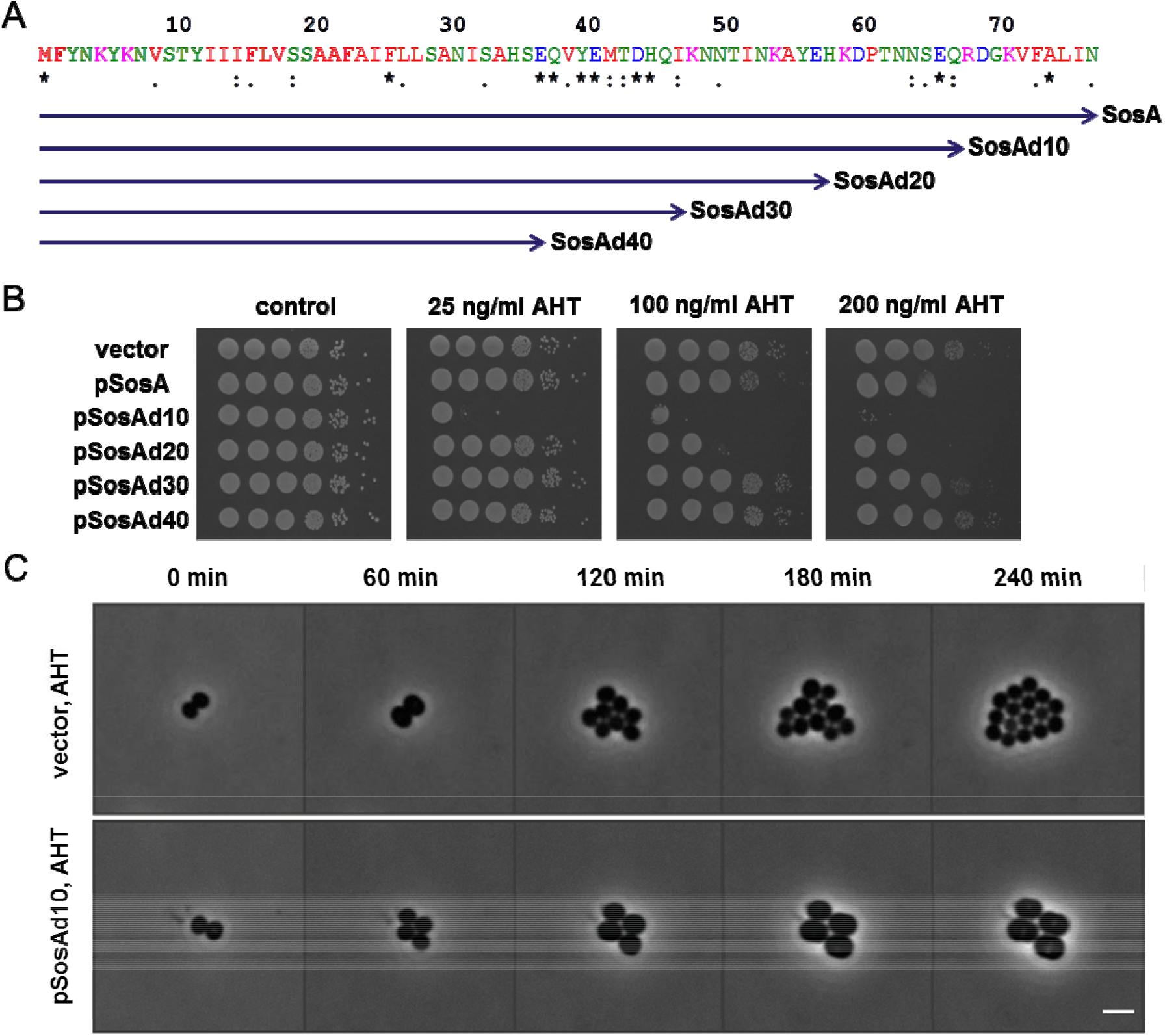
The effect of C-terminal truncations of SosA on the inhibitory activity of the protein. (A) Schematic representation of the different truncated SosA constructs. Full-length SosA is a 77-amino-acid peptide. SosAd10 lacks the extreme C-terminal 10 amino acids, while SosAd40 is SosA truncated of almost its entire extracellular C-terminal part. Indicated conserved residues (*) originate from the alignment in Figure 1. (B) Activity of the constructs was assessed inS. *aureus*RN4220 and compared to the vector control (vector). Cells were grown exponentially to an OD_600_ of 0.5, serially 10-fold diluted, and plated on TSA (control) or TSA plus inducer (AHT) at indicated increasing concentrations followed by incubation overnight at 37°C. (C) Visualization of the drastic cell size increase and the reduction in cell number of JE2/pRAB12-lacZ (control) and JE2/pRAB12-*sosAd10* in presence of 200 ng/mL of AHT by time-lapse phase-contrast microscopy at 37°C. Scale bar represents 2 μm.

**Fig 6.**
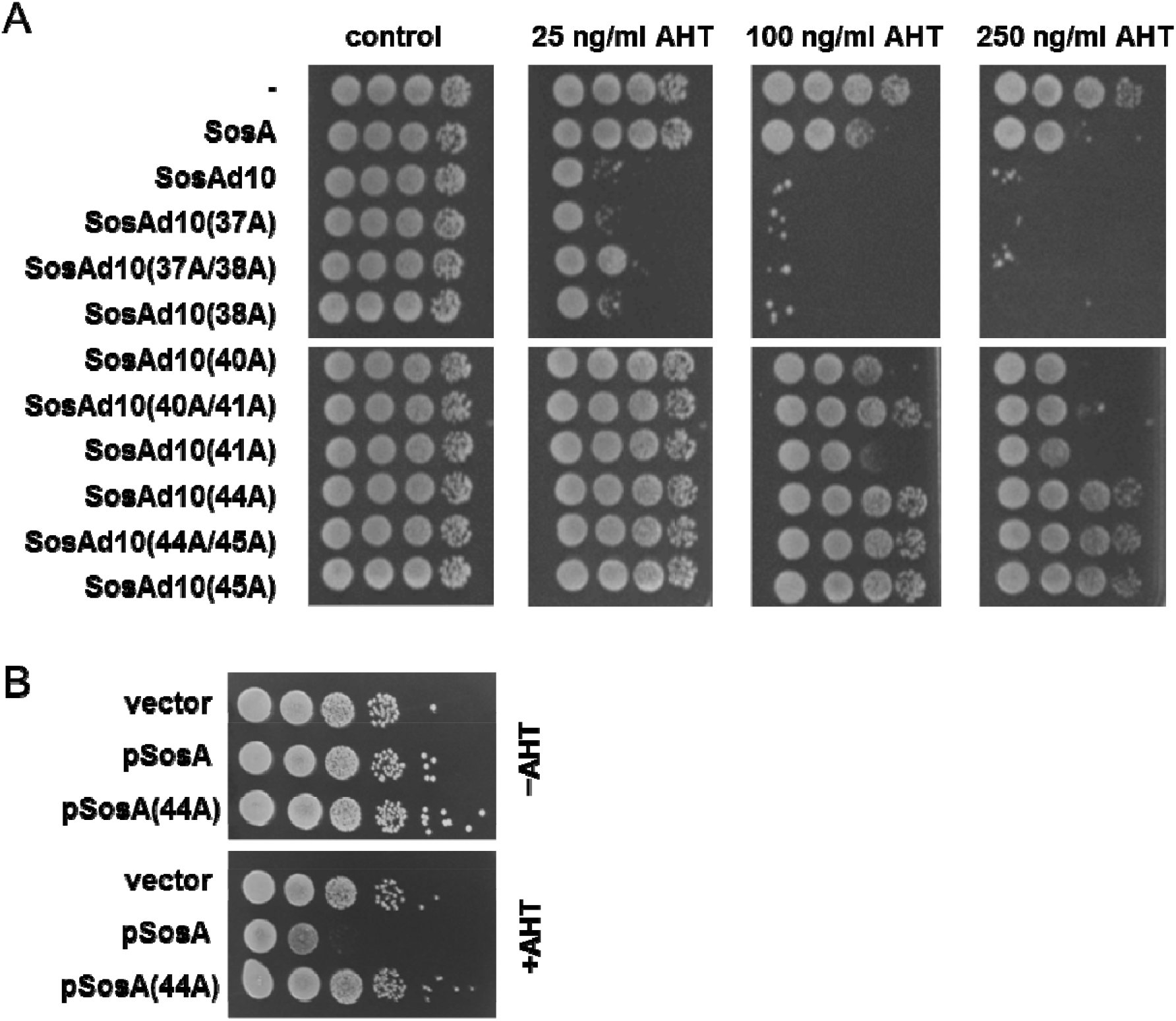
(A) Assessment of the activity of SosAd10 variants with point mutations (alanine substitutions) at conserved residues (37/38, 40/41, and 44/45) within the C-terminal part. Activity of the constructs was assessed in *S. aureus* JE2 and compared to the vector control (-), SosA, and SosAd10. Cells were grown exponentially to an OD_600_ of 0.5, serially 10-fold diluted, and plated on TSA (control) or TSA plus inducer (AHT) at increasing concentrations followed by incubation overnight at 37°C. (B) Comparison of the activity of the point mutation protein SosA(44A) with wildtype SosA in *S. aureus* JE2. Cells were grown as above and plated on TSA or TSA plus 200 ng/ml AHT.

### A CtpA protease mutant is hypersusceptible to SosA expression

We reasoned that one explanation for the severe impact of the 10-AA-truncated SosA on cell division could be that the extreme C-terminus of SosA serves as an endogenous signal for proteolytic degradation, and its removal allows the mutant protein to escape degradation and accumulate. Indeed, we observed that the cellular accumulation of the truncated protein was higher than for the wildtype protein following one hour of ectopic expression (Fig. S2E). In parallel, we observed the same pattern for cells expressing the 10-AA-truncated SosA variant with the 44A mutation that abolishes the cell division inhibitory activity suggesting that the phenotype of the mutation is not related to instability of the SosA variant (Fig. S2E). To identify a membrane localized protease that may be responsible for SosA turn-over we searched the Nebraska Transposon Mutant Library (40) forS. *aureus* protease mutant strains, and as a potential candidate, we identified the carboxyl-terminal protease A, CtpA. The *S. aureus* CtpA protein is located at the cell-membrane/cell-wall fraction and is involved in stress tolerance and virulence in a mouse model of infection (41). Importantly, overproduction of the wildtype SosA protein in cells lacking *ctpA* completely abolished cell viability, similarly to what was observed when the 10-AA-truncated SosA protein was expressed in wildtype *S. aureus* cells (Fig. 7A). In agreement with this result, we found that overproduction of SosA and CtpA in the *ctpA* mutant background alleviated the detrimental effect of SosA and that the degree of complementation depended on the relative expressional levels of the two proteins, where higher expression level of SosA was only partially complemented by CtpA expression (Fig. S3). Further substantiating the negative regulatory role played by CtpA with respect to SosA, we noted that the *ctpA* mutant was substantially more sensitive to MMC (measured by plating efficiency) than wildtype *S. aureus*, but only when encoding a functional *sosA* gene (Fig. 7B). Additionally, upon MMC-induced DNA damage, higher levels of the SosA protein accumulated in the *ctpA* mutant when compared to the wildtype (Fig. 7C). Importantly, parallel experiments were recently reported showing that CtpA in *B. subtilis* is one of two proteolytic regulators of the cell-division inhibitor YneA, with CtpA displaying direct proteolytic activity against the target in vitro. (42) Altogether, based on these results, we propose that the extracellular C-terminus of SosA is a substrate for CtpA proteolytic activity and that proteolysis relieves SosA-mediated cell-division inhibition allowing growth to resume once DNA stress has ceased.

**Fig 7.**
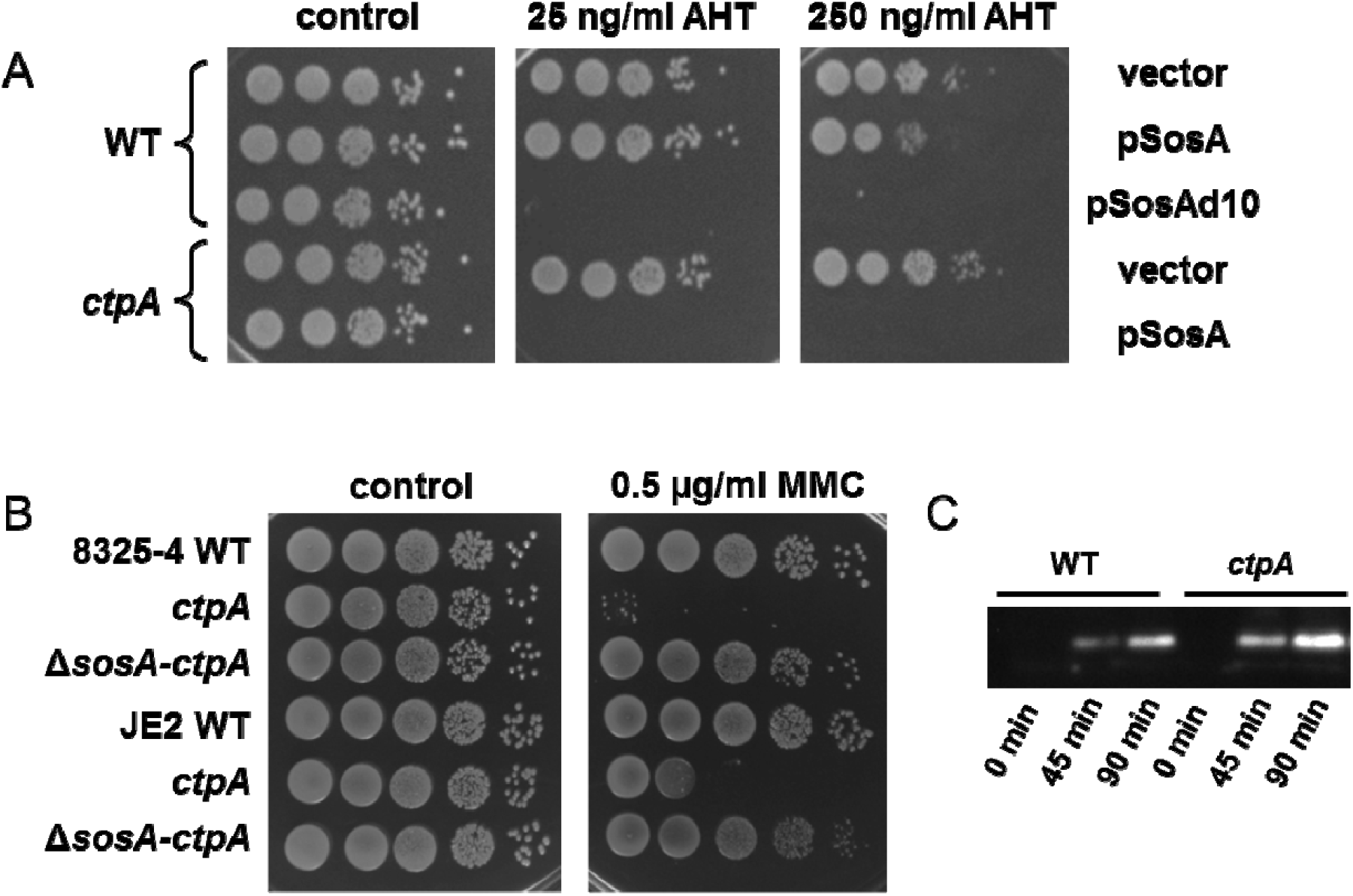
CtpA is a possible negative regulator of SosA. (A) Hypersusceptibility of an *S. aureus* JE2 *ctpA*mutant to SosA-mediated growth inhibition. Expression of *sosA* (pSosA) in wildtype *S. aureus* JE2 (WT) and the corresponding JE2-*ctpA* mutant (*ctpA*) were compared and referenced to the vector control (vector) and expression of the hyperactive SosAd10 variant (pSosAd10) in the wildtype. Cells were grown exponentially to an OD_600_ of 0.5, serially 10-fold diluted, and plated on TSA (Control) or TSA plus inducer (AHT) at indicated concentrations. The plates were incubated overnight at 37°C and imaged. (B) Assessment of mitomycin C susceptibility of a *ctpA* mutant by comparison of plating efficiency of *S. aureus* 8325-4 and JE2 wildtype, *ctpA*, and Δ*sosA-ctpA* in presence of 0.5 μg/ml MMC. Cells were grown exponentially to an OD_600_ of 0.5 and serially 10-fold diluted before plating and incubation at 37°C overnight. (C) Western blot of accumulated SosA in *S. aureus* 8325-4 wildtype (WT) and the *ctpA* mutant at indicated time points after the addition of 1 μg/ml of MMC to exponentially growing cells.

### SosA is a late-stage cell-division inhibitor

To pinpoint at which step of the cell-division process the SosA-mediated inhibition is taking place, we employed fluorescence microscopy to monitor the cellular localization of FtsZ and EzrA, crucial early-stage cell-division proteins in *S. aureus*, upon induction of SosA overproduction. Neither the *sosA* overexpression plasmid in non-inducing conditions (Fig. 8A and B) nor the control vector in non-inducing or inducing conditions (Fig. S4A and B) changed the cell morphology or protein localization. As expected, SosA overexpression caused cell swelling but also it did not cause de-localization of the cell-division initiator FtsZ (Fig. 8A) or affected the placement of the membrane-bound division protein EzrA (Fig. 8B), which is an early recruited protein connecting the cytoplasmic division components with the peptidoglycan-synthesizing membrane-bound complex (43). What was evident, however, was that whereas wildtype cells are heterogeneous with respect to FtsZ localization due to their presence in different phases of cell division, *sosA*-expressing cells appear highly synchronized with FtsZ preferentially showing septal localization (compare uninduced and induced cells in Fig. 8A)

**Fig 8.**
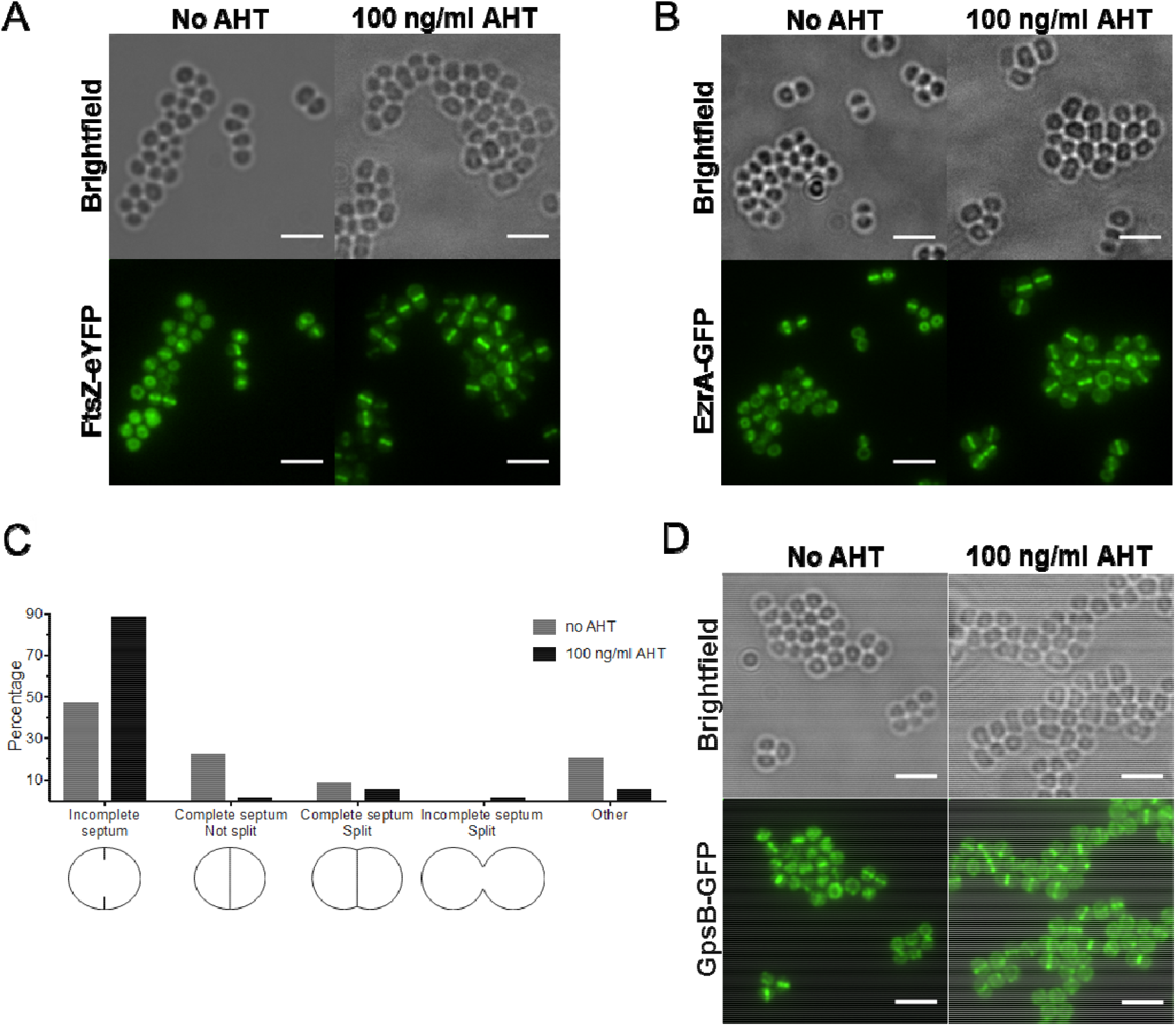
SosA does not impair the localization of cell-division proteins and septum formation initiation. (A) Localization of FtsZ-eYFP in SJF4694 (JE2 pSosA pCQ11-FtsZ-eYFP) grown in the absence and presence of 100 ng/ml of AHT for 45 min. (B) EzrA-GFP localization in SJF4697 (JE2 pSosA *ezrA-gfp*+) grown in the absence or presence of 100 ng/ml AHT for 45 min. (C) Percentages of JE2 pSosA cells exhibiting incomplete, complete, complete split, or incomplete split septa (n = 573 for no AHT, n = 607 for 100 ng/ml of AHT; organisms either isolated or co-adhered in pairs; other represents staining in indistinct shape) after incubation with or without AHT for 45 min. (D) GpsB-GFP localization in SJF700 (JE2 pSosA *gpsB-gfp*+) grown in the absence or presence of 100 ng/ml of AHT for 45 min. All fluorescence images are average intensity projections. All scale bars represent 3 um.

To further explore the cellular consequences of SosA-mediated cell-division inhibition, we examined the phenotypes of JE2/pSosA cells that were labeled for five minutes with HADA, marking regions of nascent peptidoglycan synthesis, and grown in the absence or presence of SosA induction (Fig. S4C). While in the absence of AHT, 47% and 23% of cells, respectively, showed a ring (incomplete septum) or line (complete septum) of nascent peptidoglycan synthesis, there was accumulation (89%) of cells with incomplete septa and a severe drop in cells with complete septa (1%) when SosA was overexpressed (Fig. 8C). Additionally, 1% of cells had an ‘hourglass’ shape after incubation with AHT, indicative of non-productive, premature splitting taking place prior to septum completion. Next, whole cell walls of cells overproducing SosA were labeled with a fluorescent NHS ester and examined by 3D-structured illumination microscopy (3D-SIM). Microscopy visualization revealed that most cells had only signs of septation — a so-called ‘piecrust’ (19) (Fig. S4Ei) — and showed that in an hourglass-like cell, there was a gap in the septal peptidoglycan (Fig. S4Eii). The fact that SosA does not affect localization of early cell-division proteins, FtsZ and EzrA, and the cell population becomes synchronized to a particular stage of the cell cycle (phase P2 [33]) shows that SosA does not hinder the initiation of septum formation but blocks the progression of septum completion — a phenotype similar to *S. aureus* DivlB-depleted cells (44). Although DivIB has been shown to be dispensable for FtsZ and EzrA localization to the septum and piecrust formation, its absence results in inhibition of septum progression and completion and in the mislocalization of GpsB (44), a late cell-division protein (44, 45). However, SosA overproduction did not impair the recruitment of GpsB to the mid-cell (Fig. 8D) even though visible cell enlargement occurred in this reporter strain (Fig. 8D) and not in the vector control (Fig. S4D). Altogether, these results indicate that SosA acts on (a) later cell-division component(s) than FtsZ, EzrA, and GpsB, and we conclude that SosA causes a characteristic halt in cell division by preventing progression through the morphological division stage P2, i.e. septum plate completion.

### Possible divisome interaction partners for SosA

No direct interaction has been observed between the gram-positive cell-division inhibitors and FtsZ, the scaffold for cell-division components assembly at the division site. Though poorly characterized mechanistically, the gram-positive inhibitors may suppress cell division at later stages, potentially by interacting with Ftsl/FtsQ. (for *M. tuberculosis* ChiZ) as suggested by bacterial two-hybrid analysis (36) or via delocalization of FtsL and/or DivIC (for *B. subtilis* YneA) which was only supported by indirect experimental data (46). By employing a bacterial two-hybrid system approach that reports protein-protein interactions, we sought to identify potential *in vivo* interaction candidates for SosA based on cell-division proteins from *S. aureus* (43). This analysis indicated that PBP1 (penicillin-binding protein 1), DivIC, and PBP3 are possible targets for the activity of SosA (Fig. 9A). PBP1 is a homolog of *E. coli* Ftsl (47) and an essential component of the divisome that, in *S. aureus*, is considered to play a role in septum formation and cell separation, with only the latter activity relying on transpeptidase (TP) activity of the enzyme (48). DivIC is part of an essential tripartite complex, DivlB-FtsL-DivIC (FtsQLB in *E. coli*), considered part of the second recruitment to the divisome (45, 49). Although displaying no enzymatic activity these proteins are mutually dependent on each other for stability as well being required for assembly of the final synthetic division machinery (49, 50). PBP3 is non-essential in *S. aureus* (51), but otherwise its localization and function is unknown (20). If either of these proteins are indeed targets for SosA, we predicted that ectopic expression of *pbpA* (encoding PBP1), *divIC*, or *pbpC* (encoding PBP3) would mitigate the effect of concomitant SosA overexpression by increasing the level of target protein. While no effect was observed upon DivIC and PBP3 co-overexpression, overproduction of PBP1 along with SosA was manifested as a synergistically lethal phenotype with to considerable growth impairment (Fig. 9B). Further, we found that this phenotype was apparent even at low inducer concentrations, where the effect of SosA overproduction alone is negligible, and that it was independent on the transpeptidase activity of PBP1 as overexpression of a mutant variant of PBP1 lacking this activity together with SosA still eliminated growth (Fig. S5A). The phenotype is, however, dependent on the cell-division inhibitory activity of SosA as concomitant overproduction of the SosA(44A) mutant protein abolished the phenotype (Fig. S5B), suggesting that the lethal phenotype is not a simple artifact of ectopic overexpression of a membrane protein along with PBP1. While these data do not support a simple titration of SosA by PBP1, it does support that SosA is mechanistically linked to, or intercepts with the activity of PBP1. It is also conceivable that SosA and PBP1 act synergistically on the same target protein, the former by inhibiting its activity and the latter via disrupting the stoichiometry of the division components. Lastly, we examined the importance of the inactivating mutation (44A) of SosA as well as the stabilizing (SosAd10) and inactivating (SosAd40) truncations to the protein for interaction with PBP1 and DivIC, respectively. Interestingly, we find that the inactivating mutation compromises interaction with DivIC only, whereas both interactions are dependent on the C-terminus of SosA (Fig. 9C). This indicates that the inhibitory activity of SosA is functionally related to its interaction with DivIC.

**Fig 9.**
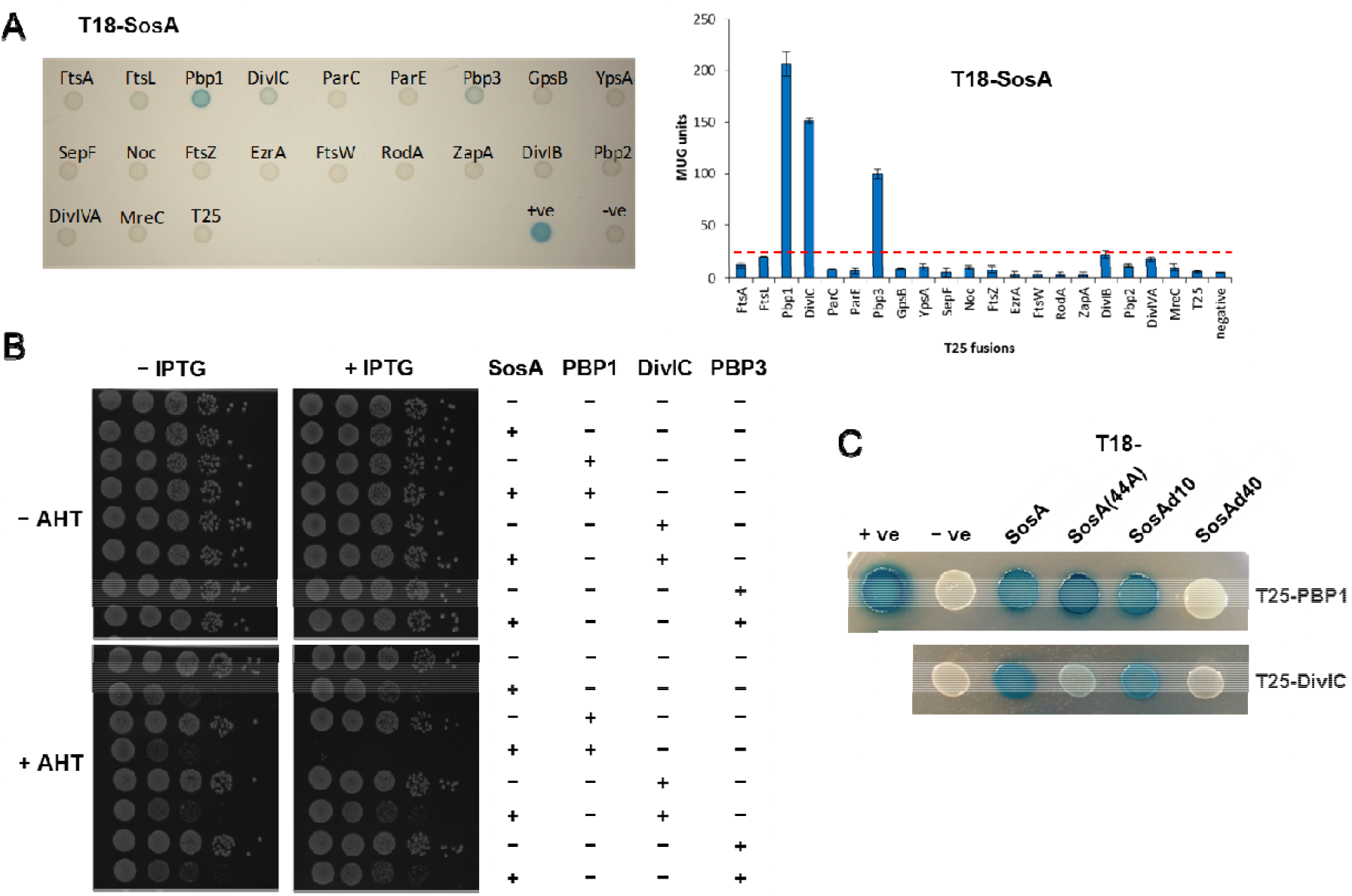
SosA interaction with the central divisome components. (A) Analysis of the protein interactions between SosA and a panel of *S. aureus* cell-division or cell wall-synthesis proteins using the bacterial two-hybrid system. T18 and T25 are domains of an adenylate cyclase enzyme, and the “T18-SosA” and “T25 fusions” labels indicate the fusions between the C-terminus of the T18 adenylate cyclase fragment to the N-terminus of SosA and the T25 fusions to the cell-division proteins. *Left*: Interactions between the *S. aureus* cell-division proteins fused to T25 and *S. aureus* SosA fused to T18, wherein 10 μl of a 1:100 dilution of overnight culture of co-transformed BTH101 were spotted onto minimal medium containing 150 μg/ml of X-gal and incubated at 30°C for 36 h. +ve, T25-zip + T18-zip; −ve, pKT25 + pUT18C. *Right*: β-Galactosidase activity of interactions between *S. aureus* T18-SosA and cell-division proteins (T25 fusions). Activity is displayed as the mean of three independent measurements of β-galactosidase activity, in MUG (4-methylumbelliferyl-β-D-galactopyranoside) units, for each co-transformant. Error bars represent the standard deviation. Positive interactions are considered to be at least four times higher than the activity level for the negative control, and this cut-off level is represented by the red line in the bar chart. (B) Assessment of the plating efficiency of *S. aureus* JE2 upon concomitant overexpression of *sosA* (pSosA, AHT-inducible) and *pbpA* (pPBP1) or *divIC* (pDivIC) or *pbpC* (pPBP3) controlled by IPTG (+). (-) denotes the vector controls. Cells were grown exponentially to an OD_600_ of 0.5, serially 10-fold diluted, and plated on TSA plates containing 100 ng/ml AHT and 0.2 mM IPTG where indicated. The plates were incubated overnight at 37°C and imaged. (C) Interactions of T18 fusions to SosA, SosA(44A), SosAd10 and SosAd40 with T25-PBP1 or T25-DivIC. 10 μl of a 1:100 dilution of overnight cultures of co-transformed BTH101 were spotted onto LB containing 150 μg/ml of X-gal and incubated at 30°C for 24 h. +ve and −ve denotes positive and negative interaction controls, respectively.

## Discussion

Based on the results reported here, we propose the following model for regulation of cell division in the prolate spheroid bacterium *S. aureus* under DNA-damaging conditions (Fig. 10). During normal growth, the expression of *sosA* and the entire SOS regulon is repressed by the LexA repressor. Under DNA damaging and SOS inducing conditions, LexA is inactivated through autocleavage, which results in the expression of *sosA*. In this process, the N-terminal autocleavage product of LexA needs to be further proteolytically processed for *sosA* expression to be fully induced (23), thus adding another layer of expressional control, suggesting tight regulation of the inhibitory protein. Upon derepression of LexA-regulated genes, SosA accumulates to levels that inhibit cell division without causing FtsZ delocalization nor preventing the assembly of downstream components, EzrA and GpsB, which are representatives of early and later cell division components. The inhibitor appears to affect the progression and completion of septum formation via interaction with essential divisome components with DivIC being one such possible target candidate. As a consequence of SosA activity, the cells appear synchronized at the stage of septum formation initiation, while an ongoing off-septal peptidoglycan synthesis leads to cell enlargement (Fig. S4C). Once conditions are favorable for continued division, *sosA* expression is repressed, the membrane protease CtpA degrades SosA, and cell division resumes (Fig. 10).

**Fig 10.**
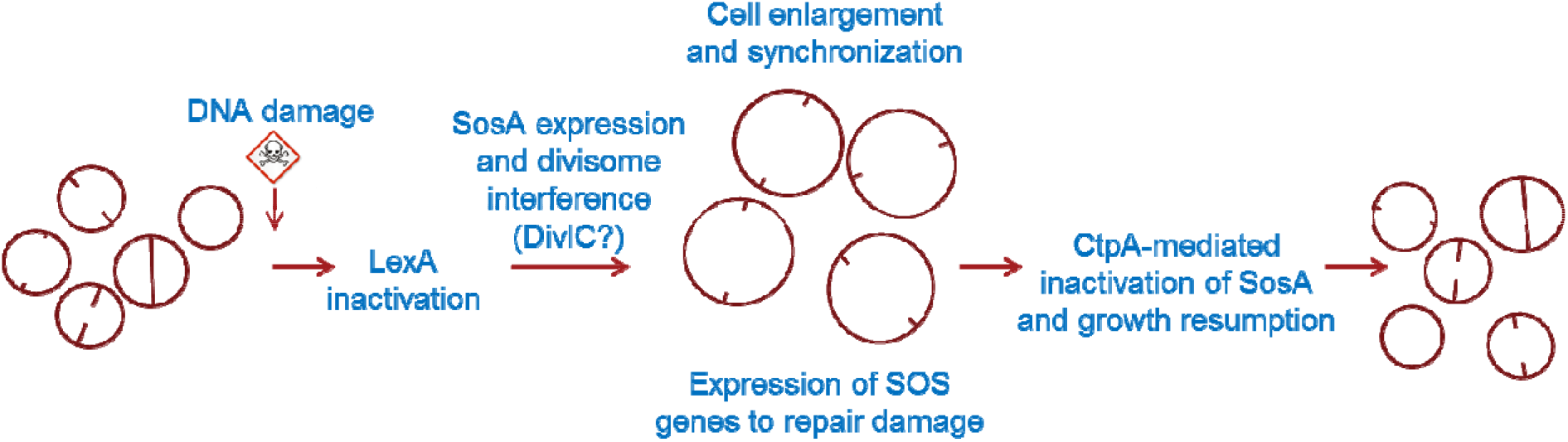
Proposed model for a regulated survival strategy for *S. aureus* upon DNA damage. As part of the SOS response, SosA is produced and, by a membrane-localized activity, corrupts normal cell-division activity, likely via an interaction with essential divisome components. In effect, cells are still able to initiate septum formation but are unable to complete it, leading to synchronization of the cell population. This provides a spatio-temporal window for paralleled SOS response-regulated DNA-repair activity. At the same time, the cell size increases due to off-septal activity of peptidoglycan-synthesis enzymes. As the SOS response diminishes, the cellular concentration of SosA is lowered, directly or indirectly, by the proteolytic activity of CtpA. At this stage, septa are allowed to complete and normal growth/division continues.

The model illuminates several remaining unanswered questions. It will be particularly important to determine if and how SosA affects the function of DivIC and by which means SosA intercepts the activity of PBP1. In this context, it should be noted that our data fit well with the finding that the genetic depletion of PBP1 causes cell enlargement and incomplete septum formation (47). At present we cannot readily explain the peculiar finding that SosA and PBP1 seem to act in a synergistically lethal manner (Fig. 9, and Fig. S5). Though it is formally possible that the two proteins together constitute a signal to halt septum formation beyond a certain point, it is plausible that our observations are due to severe changes in stoichiometry within the divisome. It is also conceivable that overexpression of PBP1 is prematurely forcing septum completion which could be counterproductive under the inhibitory activity of SosA. In fact, we find that overexpression of another essential component of the peptidoglycan synthesis machinery, the lipid II flippase MurJ, is also synthetically lethal when SosA is expressed (Fig S5C), showing that the phenotype is not exclusively related to PBP1. Our microscopy and localization data suggest that SosA acts quite specifically at a decision point between septum initiation and septum completion. In this regard, it is interesting that cell division in *S. aureus* was recently reported to be a two-step process characterized by an initial, slow FtsZ-dependent activity followed by a faster process when MurJ arrives and directs peptidoglycan synthesis to the septum (in division stage P2 where SosA expressing cells are stalled) (35). More intriguingly, MurJ recruitment was reported to be mediated by the DivlB-FtsL-DivIC complex (35), which could constitute an important molecular cue for the inhibitory activity of SosA. Along this line, FtsL and DivIC are considered unstable proteins, and in *B. subtilis* stability of DivIC is dependent on FtsL (52). Hence, we tried to co-express FtsL along with SosA and did note a partial alleviation of the inhibitory activity (Fig. S5D). We take this observation as another indication that DivIC could be a direct target of SosA, although at present we cannot exclude indirect effects. Interactions between the *S. aureus* divisome components constitute a complex web, and in a previous bacterial two-hybrid study it has been shown that PBP1 interacts with DivIC, which in turn interacts with PBP3 (43). Thus, the inhibitory activity of SosA could be multifactorial and further studies are required to delineate the exact function of the protein. Such studies should include localization studies of e.g., DivIC and MurJ as recently employed (35) but could also include mutational and suppressor studies as elegantly used to map the cell-division inhibition process in *C. crescentus* (16,17).

Analysis of the AA sequence of SosA indicated that the N-terminal part contains a TM domain while the C-terminal is located ext race 11 u la r ly. The hypothesis that the TM domain promotes localization to the cellular membrane was supported by PhoA-LacZ fusion to SosA and by various point and deletion mutants of the fusion protein (Fig. 4). The abrogation of activity by the 40-AA-truncation as well as by a single alanine substitution (D44A) suggests that this segment is essential for correct localization to the septum or, perhaps more interestingly, that the actual motif for cell-division inhibition is localized at the exterior of the membrane. In this respect, it is interesting that the single amino acid substitution affects the interaction with DivIC in the two-hybrid system (Fig. 9C). If divisome interference indeed lies within this extracellular segment of SosA, it could provide a novel paradigm for interference with cell-division beyond the transpeptidase-inhibitory activity exerted by ß-lactam antibiotics.

Endogenously triggered cell-division inhibition must be strictly regulated, as demonstrated by pioneering work showing that SulA of *E. coli* is degraded by the Lon protease (15). Similarly, YneA from *B. subtilis* was reported to be regulated by proteolysis, by an undefined mechanism, and a point mutation at its extreme C-terminus generated a stabilized yet functional variant of the cell-division inhibitor (37). In *B. megaterium*, the temporal expression of a YneA homologue has been suggested to be due to mRNA instability (26). We demonstrate that a minor truncation of the C-terminus of SosA increases cell division inhibitory activity (Fig. 5), possibly as a consequence of accumulation of the protein (Fig. S2E). This led us to speculate that the extreme C-terminus serves as a signal for proteolysis and we identified the CtpA protease-deficient strain in which SosA appeared to accumulate and had an increased detrimental effect compared to the wildtype background. Bacterial carboxy-terminal proteases are poorly characterized, particularly with respect to their substrate preferences; however, we propose that CtpA specifically regulates SOS-induced cell-division inhibition in staphylococci, likely via recognition of the C-terminus of SosA. In support of this, Burby et al. (42) recently disclosed that CtpA is involved in turn-over of YneA in *B. subtilis*. Direct biochemical evidence for such a direct activity of CtpA on SosA in *S. aureus* is needed, but a shared mechanism is highly conceivable. Of note, carboxy-terminal proteases are ubiquitous in bacterial genera, and homologs of CtpA are found in other gram-positive species, including also *L monocytogenes* which share a homolog of YneA. Hence, the CtpA-like proteases could represent functional homologs of the *E. coli* Lon protease, allowing negative regulation of membrane-localized endogenous cell-division inhibition more broadly amongst Firmicutes and beyond.

SosA may also prove to be a useful tool for scientists conducting studies of bacterial cell division, which have recently benefitted from the development of genetic tools and highly sophisticated microscopic imaging technologies. However, most often bacteria grow as an unsynchronized population with individual cells in different phases of cell division, making it challenging to order the temporal events that occur during cell-cycle progression, DNA replication, and division. Essentially, SosA is involved in halting the *S. aureus* cell cycle at a specific morphological stage, e.g. following the piecrust formation, and we anticipate that the exploitation of this function may constitute a useful tool for the generation of synchronized staphylococcal cells for experimental purposes.

Apart from its basic biological interest, the bacterial cell-division process may constitute an unexhausted source of novel therapeutic targets (1,2). We believe that SosA could be a natural scaffold, and that further studies on this protein can provide new therapeutically accessible targets that would interfere with an essential process in *S. aureus* and other staphylococci.

## Materials and Methods

### Bacterial strains and plasmids used in this work

The bacterial strains and plasmids used in this study are listed in Table S1. *E. coli* strains were grown in Luria-Bertani medium (Oxoid) or on LB agar (Oxoid). *S. aureus* strains were grown in Tryptic Soy Broth (TSB) or on Tryptic Soy Agar (TSA) (both Oxoid). Ampicillin (100 μg/ml), chloramphenicol (10 μg/ml), or erythromycin (5 μg/ml) were added when appropriate. Anhydrotetracycline (AHT) (Sigma-Aldrich) and Isopropyl β-D-1-thiogalactopyranoside (IPTG) (Thermo Scientific) were used as inducers for protein expression.

### Construction of strains and plasmids

All oligonucleotides used are listed in Table S1. *S. aureus*8325-4ΔsosA is a clean deletion of *sosA* (SAOUHSC_01334) in strain 8325-4 obtained by allelic replacement using the temperature-sensitive shuttle vector pIMAY (53). 1 kb regions up- and downstream of *sosA* were PCR amplified (Phusion Hot Start II DNA Polymerase, Thermo Scientific) using primer pairs Up-sosA_fw-Kpnl/Up-sosA_rev and Dw-sosA_fw/Dw-sosA_rev-SacI, respectively, and subsequently joined in a spliced overhang PCR using Up-sosA_fw-Kpnl/Dw-sosA_rev-Sacl. The resulting deletion fragment was cloned into pIMAY via Kpnl/Sacl, generating pIMAY-ΔsosA purified from *E. coli* DC10B and transformed into *S. aureus* subsequently maintained at 28°C. Chromosomal integration of the plasmid was performed at 37°C under selective conditions (chloramphenicol) followed by passage at 28°C without antibiotic selection and final plating on TSA containing 500 ng/ml of AHT for plasmid counterselection. Colonies were replica-plated to select for sensitivity towards chloramphenicol and successful allelic exchange were screened for by PCR amplification using primer pairs Ctrl_dsosA_F/Ctrl_dsosA_R positioned outside *sosA* and Fwd_MCS/Rev_MCS targeting the vector, respectively. The allelic replacement procedure was identical for *S. aureus* JE2 to create JE2ΔsosA.

The chromosomal transposon insertion mutation in *ctpA* conferring erythromycin resistance was obtained from the Nebraska Transposon Mutant Library (NTML, NE847) (40) and was moved by transduction (phage (ϕll) to *S. aureus* JE2, JE2Δ*sosA S. aureus* 8325-4 and 8325-4Δ*sosA* resulting in JE2-*ctpA*, JE2Δ*sosA-ctpA*, 8325-4-*ctpA*, and 8325-4Δ*sosA-ctpA*, respectively. JE2-*ctpA*(–erm), an erythromycin-sensitive derivative, was generated by elimination of the transposon-encoded *ermB*in JE2-*ctpA* by allelic exchange using the pTnT vector using temperature-mediated chromosomal integration and secY-mediated counterselection as described before (54).

All plasmid constructs mentioned below were cloned in E. coli IM08B (55) from where they were transformed into *S. aureus* strains. For complementation of the *sosA* knockout, the *sosA* gene with its native promoter was PCR amplified from strain 8325-4 using primers Up-sosA-promo_Sall and Dw-sosA_EcoRI and cloned into designated restriction sites in plasmid pJC1112 (56). The resulting plasmid (pJC1112-sosA) was transformed into the integration-proficient strain RN9011, creating RN9011-sosA-compl. when selected with erythromycin. The chromosomally integrated plasmid was transduced into 8325-4Δ*sos*A resulting in 8325-4Δ*sosA*-compl. ForsosA expression the *sosA* gene including its predicted ribosomal binding site was cloned into the BglII and EcoRI sites of pRAB12-*lacZ* (57) using primers Up-*sosA*_BglII/Dw-sosA_EcoRI, at the same time eliminating the *lacZ* reporter gene. The resulting plasmid was called pSosA. Generation of plasmids encoding C-terminally truncated variants of SosA was obtained by PCR amplification of DNA fragments using Up-sosA_BglII and downstream primers Dw-sosA(d10)_EcoRI to Dw-sosA(d40)_EcoRI, all equipped with premature stop codons. The PCR products were ligated into pRAB12-*lacZ* using Bglll and EcoRI cut sites to create pSosAd10, pSosAd20, pSosAd30 and pSosAd40. Single and double amino acid substitutions (alanine) in the SosAd10 protein were obtained by cloning commercially synthesized DNA fragments (Twist Bioscience) into pRAB12-*lacZ* using the same restriction sites. The mutation at AA position 44 (D to A) was introduced to the full length SosA protein via extension PCR by first using primers Up-sosA_Bglll/SosA_R-long on the SosAd10(44A) template, then using that product as template for amplification with primers Up-sosA_BglII/Dw-sosA_EcoRI followed by cloning into pRAB12-*lacZ*, thereby generating pSosA(44A). Expression plasmids pPBPl, pDivIC, pPBP3, pFtsL, pMurJ, and pCtpA were constructed by cloning the pbpA*divIC, pbpC, ftsL, murJ*, and *ctpA* genes behind the P_spac_ promoter in pSK9067 (58) using primer pairs pbpA_F-Sall/pbpA_R-EcoRI, divlC-F_Sall/divlC-R_EcoRI, pbpC-F_Sall/pbpC-R_EcoRI, ftsL-F_Sall/ftsL-R_Aatll, murJ-F_Sall/murJ-R_Aatll, and ctpA_F-Sall/ctpA_R-EcoRI, respectively. A plasmid for *pbpA\** expression was constructed by use of the same primer pair as for *pbpA* but using pMAD-PBPl* (59) as a template. For membrane topology analysis using pKTop (39), *sosA, sosAd10, sosAd10(44A)* and *sosAd40* were PCR amplified from respective expression constructs using primer SosA_F-BamHI and primers SosA_R-Kpnl, SosAd10_R-Kpnl or SosAd40_R-Kpnl, respectively, and cloned in frame in front of the *phoA-lacZ* chimeric gene using BamHI/Kpnl restriction sites.

In order to construct JE2 strains producing a fluorescent fusion of FtsZ, EzrA or GpsB in the presence of the *sosA* expression plasmid, JE2 pSosA was transduced with a lysate from SH4665 (SH1000 pCQ11-FtsZ-eYFP), JGL227 (SH1000 *ezrA-gfp*+) or JGL228 (SH1000*gpsB-gfp*+), resulting in SJF4694 (JE2 pRAB12-*lacZ* pCQ11-FtsZ-eYFP), SJF4696 (JE2 pRAB12-*lacZ ezrA-gfp*+) and SJF4699 (JE2 pRAB12-*lacZ gpsB-gfp*+), respectively. Control strains SJF4693 (JE2 pRAB12-*lacZ* pCQ11-FtsZ-eYFP), SJF4696 (JE2 pRAB12-*lacZ ezrA-gfp*+) and SJF4699 (JE2 pRAB12-*lacZ gpsB-gfp*+) were constructed by a phage transduction of JE2 pRAB12–/ocZ with lysates from SH4665 (SH1000 pCQ11-FtsZ-eYFP), JGL227 (SH1000 *ezrA-gfp*+) or JGL228 (SH1000 *gpsB-gfp+)*, respectively. To induce FtsZ-eYFP production in SJF4693 and SJF4694, cells were grown in the presence of 50 μM IPTG.

In order to screen for interaction of SosA with cell-division/cell wall synthesis proteins of *S. aureus, sosA* was amplified from *S. aureus* SH1000 genomic DNA using primers ALB133 and ALB134, and ligated into pUT18C using BamHI/EcoRI cut sites, resulting in pT18-SosA. T18 fusions with SosAd10 and SosAd40 were generated by combining ALB133 with respective truncation primers (Dw-sosA(d10)_EcoRI and Dw-sosA(d40)_EcoRI). The pT18-SosA(44A) plasmid was obtained as above but by use of pSosA(44A) as PCR amplification template. All fusion constructs to T25 and control vectors were generated previously (43, 60-62, Table SI).

### Determination of OD_600_ and CFU after MMC treatment

Strains were grown overnight on TSA plates and used for inoculating TSB and allowed to grow to OD_600_ = 0.05 when Mitomycin C (MMC from *5treptomyces caespitosus*, Sigma-Aldrich) was added. Cell density was monitored by OD_600_ measurements at intervals onwards and samples were withdrawn to determine culture colony forming units by serial dilution in 0.9% w/v NaCI and plating on TSA.

### Flow cytometry

Cell size distributions of cultures were arbitrarily quantified by flow cytometry using the forward scatter signal (FSC-A) acquired on a BD Accuri™ C6 flow cytometer (BD Biosciences). Cell samples were diluted in 0.9% w/v NaCI to an approximate density of 10^6^ cells/ml and sonicated briefly to allow acquisition of scatter signal from single cells predominantly. Sonication was performed with a Bandelin sonopuls HD2070/UW2070 apparatus (Bandelin electronics, Germany) fitted with the MS 73 probe. Ten pulses of 500 msec were given at 50% power. All flow cytometry experiments were independently repeated at least twice with similar results.

### Plating assays

Spot dilution was used to evaluate plating efficiency of strains carrying various plasmid constructs. Strains were grown exponentially until an approximate OD_600_ of 0.5 under plasmid selective conditions. Strains were then ten-fold serially diluted in 0.9% w/v NaCI and positioned as 10 μl spots on TSA containing selective antibiotics with/without inducer at indicated concentrations and incubated at 37°C overnight. All spot plating assays were independently repeated with similar results.

#### Labelling *S. aureus* with HADA

Cells grown to mid-exponential phase (OD_600_ ~ 0.5) were incubated with 500 μM HADA at 37°C for 5 min. Cells were then washed by centrifugation and resuspension in PBS.

#### Labelling *S. aureus* with NHS ester

Labelling with NHS ester was performed as described before (34). Briefly, cells grown to midexponential phase (OD_600_ ~0.5) were collected by centrifugation and growth medium was discarded. Cells were resuspended in PBS containing 8 μg ml^-1^ Alexa Fluor 647 NHS ester (Invitrogen) and incubated at room temperature for 5 min. Cells were washed by centrifugation and resuspension in PBS.

#### Fixing

Cells were fixed by incubation in 4% (w/v) paraformaldehyde at room temperature for 30 min.

#### Widefield Epifluorescence Microscopy

Fixed cells were dried onto a poly-L-Lysine coated slide and mounted in PBS. Imaging was performed using either a Nikon Ti Inverted microscope fitted with a Lumencor Spectra X light engine or a v4 DeltaVision OMX 3D-SIM system (Applied Precision, GE Healthcare, Issaquah, USA).

#### 3D Structured Illumination Microscopy

A high precision cover slip (High-precision, No.1.5H, 22×22mm, 170±5 μm, Marienfeld) was cleaned by sonicating in 1 M KOH for 15 min at room temperature. The coverslip was washed with water and incubated in 0.01% (w/v) poly-L-lysine solution (Sigma) for 30 min at room temperature. The coverslip was rinsed with water and dried with nitrogen. Fixed cells were dried onto the poly-L-Lysine coated cover slip and mounted on a slide with SlowFade Diamond (Invitrogen). 3D SIM visualisation was performed using a v4 DeltaVision OMX 3D-SIM system (Applied Precision, GE Healthcare, Issaquah, USA) equipped with a Plan Apo 60x, 1.42 NA oil objective, using 1.514 immersion oil, a 642 nm laser and a standard excitation/emission filter set (683/40). For each z-section, a sample was imaged in 5 phase shifts and 3 angles. The z-sections were 0.125 nm in depth. Raw data were reconstructed with the Softworx software (GE Healthcare) using OTFs optimized for the specific wavelength and oil used.

### Time-lapse microscopy

Strains were grown overnight in TSB medium at 37°C, then diluted 100 times in fresh TSB and grown until OD = 0.1. Cells were washed once with fresh TSB and spotted onto TSB-acrylamide (10%) pads previously incubated for 2h in TSB medium supplemented, when appropriate, with 0.04 μg/ml. mitomycin C (MMC) or 200 ng/mL anhydrotetracycline (AHT). Pads were placed into a Gene frame (Thermo Fisher Scientific) and sealed with a cover glass. Phase-contrast images were acquired on a DV Elite microscope (GE healthcare) equipped with a sCMOS (PCO) camera and a 100x oil-immersion objective. Images were acquired with 200 ms exposure time every 4 minutes for at least 6 h at 37°C using Softworx (Applied Precision) software. Images were analyzed using Fiji (http://fiii.sc).

### Western blot analysis

Purified rabbit anti-SosA antibody was obtained via GenScript Biotech (Netherlands). A custom peptide, CEHKDPTNNSEQRDGA, comprising AAs 57-70 of SosA was used for immunization. Bacterial cells were pelleted and frozen immediately at −80°C before being resuspended in PBS containing cOmplete™ Mini Protease Inhibitor Cocktail (Sigma-Aldrich) and lysed by bead-beating (Fastprep-24™, MP Biomedicals). The protein concentration of the lysates was measured using the Qubit™ Protein Assay Kit (ThermoFisher Scientific) and samples were normalized to equal amounts of protein, 20 μg of total protein was separated on NuPAGE^®^ 4-12% Bis-Tris gels using MES buffer and the XCell sure-lock mini-cell system (ThermoFisher Scientific), transferred to a polyvinylidene difluoride membrane(PVDF) and probed with primary antibody diluted 1:1000. Bound antibody was detected with the WesternBreeze^®^ Chemiluminescent Kit, anti-rabbit according to the instructions from the manufacturer (ThermoFisher Scientific). All western blots were independently repeated with similar results. Uncropped versions of blots are displayed in Fig S6.

### Bacterial two-hybrid assay

Interactions between SosA and proteins involved in cell division and cell wall synthesis of *S. aureus* were tested using *E.coli* BTH101 (Δ*cyaA*) cotransformed with pT18-SosA and a plasmid carrying a fusion of T25 with a cell-division or cell wall synthesis protein on minimal medium as previously described (43). Mutant and truncated variants of SosA were tested specifically for interaction with PBP1 and DivIC in comparison to the interaction observed with the wildtype protein.

## Acknowledgements

This work was supported by grants from the Danish Council for Independent Research – Technology and Production (1337-00129 and 1335-00772) to M.S.B., by the Danish National Research Foundation (DNRF120) to H.I., and by the Medical Research Council (MR/K015753/1 to the Wolfson Light Microscopy Facility at the University of Sheffield and MR/N002679/1). We would like to thank Prof Simon Jones (University of Sheffield) for the HADA dye.

M.S.B., K.W., P.K., M.T.C., S.J.F., and H.l. conceived and designed the study. Experiments were performed by M.S.B., K.W., C.G., and A.L.B. M.S.B. K.W., C.G., G.L., D.F., J.-W.V., S.J.F., and H.l. contributed to analysis of data and drafting of the manuscript. All authors read and approved the final manuscript.

## Supplemental Material

6 supplementary figures, 1 supplementary table, 3 supplementary movies.

### Legends for supplemental material

**Fig.** SI Percent identity matrix for staphylococcal SosA proteins. Pairwise identity scores for the proteins included in the alignment in Figure 1 were obtained by the Clustal 2.1 algorithm.

**Fig. S2** (A) Detection by western blot of DNA-damage induced expression of SosA. SosA being detected in wildtype *S. aureus* 8325-4 and the complemented strain, while being absent in the *sosA* deletion mutant following 1 h of MMC treatment (1 μg/ml). Also displayed is the time-dependent expression of SosA in *S. aureus* JE2 post MMC addition (1 μg/ml) in comparison to the corresponding *sosA* deletion mutant. (B and C) Phenotypic complementation of thesosA deletion mutant. The phenotypes of *S. aureus* 8325-4ΔsosA were restored back to wildtype by chromosomal integration of a copy of the sosA gene under its native promoter (*S. aureus* 8325-4ΔsosA-compl.) when evaluated for changes in cell density and viability (B) and cell size assessed by flow cytometry (C) during challenge with MMC (1 μg/ml). Mean FSC-A values are indicated below histograms. Cells were grown exponentially prior to addition of MMC at an OD_600_ of 0.05. (D) Effect of different truncated SosA variants on *S. aureus* cell size. Evaluation of cell size distribution was performed by flow cytometry (FSC-A) of *S. aureus* RN4220 containing expression plasmids encoding full length SosA or C-terminally truncated variants of the protein. Mean FSC-A values are indicated below histograms. Cells were grown exponentially prior to induction with 100 ng/ml of AHT and analyzed at indicated time points. (E) Western blot of accumulation in *S. aureus*JE2 of SosA, SosAd10, SosA(44A), and SosAd10(44A) expressed from respective plasmid constructs for 1 h with 200 ng/ml AHT.

**Fig. S3** Hypersusceptibility of an *S. aureus* JE2 *ctpA* mutant to SosA-mediated growth inhibition. Plating efficiency of the *S. aureus* JE2-ctpA(-erm) mutant transformed with both SosA and CtpA expression plasmids (+) or vector controls (-) at different inducer concentrations; AHT for SosA and IPTG for CtpA. Cells were grown exponentially to an OD_600_ of 0.5, serially 10-fold diluted, and plated on TSA plus indicated inducer concentrations followed by incubation overnight at 37°C before imaging.

**Fig. S4** SosA halts septum completion. Localization of (A) FtsZ-eYFP in SJF4693 (JE2 pRAB12 -*lacZ*pCQ11-FtsZ-eYFP), (B) EzrA-GFP in SJF4696 (JE2 pRAB12-*lacZezrA-gfp*+) and (D) GpsB-GFP in SJF4699 (JE2 pRAB12-*lacZ gpsB-gfp*+) grown in the absence and presence of 100 ng/ml of AHT for 45 min. Fluorescence images are average intensity projections. Scale bars represents 3 μm. (C) Fluorescence microscopy images of JE2/pSosA grown in the absence or presence of 100 ng/ml of AHT for 45 min and labeled with HADA for 5 min. Images are average intensity projections. Scale bars represents 3 μm. (E) 3D-SIM Z-stack images of JE2/pSosA grown with 100 ng/ml of AHT for 45 min and labelled with Alexa Fluor 647 NHS ester, (i) A cell with initiated septum formation and (ii) a cell splitting into daughter cells without finishing a septal disc. Scale bars represents 0.5 μm.

**Fig. S5** Assessment of the plating efficiency of *S. aureus* strain JE2 upon expression of sosA (pSosA, AHT-inducible) and concomitant overexpression of PBP1 (*pbpA*) or a PBP1 transpeptidase mutant PBP1* (*pbpA*\*) (A), PBP1 (*pbpA*) and in comparison to SosA(44A) (B), MurJ (C), and FtsL (D). Inducer concentrations for SosA (and SosA(44A)) were 50 ng/ml (panels A-C) and 150 ng/ml AHT (panel D). Co-expression proteins were induced by 0.2 mM IPTG. (+) denotes protein encoding vectors, (-) denotes empty vector controls. In all panels, cells were grown exponentially to an OD_600_ of 0.5, serially 10-fold diluted, and plated on TSA with/without inducers followed by incubation overnight at 37°C before imaging.

**Fig. S6** Uncropped western blots relating to Figure 7 and Figure S2.

**Movie SI.** Time-lapse microscopy of *S. aureus* JE2 WT and sosA mutant upon MMC exposure.

**Movie S2.** Time-lapse microscopy of *S. aureus* JE2 cells overexpressing SosA.

**Movie S3.** Time-lapse microscopy of *S. aureus* JE2 cells overexpressing SosAd10.

## References

1. Sass P, Brötz-Oesterhelt H. 2013. Bacterial cell division as a target for new antibiotics. Curr Opin Microbiol 16:522–530.

2. Lock RL, Harry EJ. 2008. Cell-division inhibitors: new insights for future antibiotics. Nat Rev Drug Discov 7:324–338.

3. Robinson A, Causer RJ, Dixon NE. 2012. Architecture and conservation of the bacterial DNA replication machinery, an underexploited drug target. Curr Drug Targets 13:352–372.

4. Adams DW, Errington J. 2009. Bacterial cell division: assembly, maintenance and disassembly of the Z ring. Nat Rev Microbiol 7:642–653.

5. Baharoglu Z, Mazel D. 2014. SOS, the formidable strategy of bacteria against aggressions. FEMS Microbiol Rev 38:1126–1145.

6. Kreuzer KN. 2013. DNA damage responses in prokaryotes: regulating gene expression, modulating growth patterns, and manipulating replication forks. Cold Spring Harb Perspect Biol 5:a012674.

7. Kelley WL. 2006. Lex marks the spot: the virulent side of SOS and a closer look at the LexA regulon. Mol Microbiol 62:1228–1238.

8. Huisman O, D’Ari R. 1981. An inducible DNA replication-cell division coupling mechanism in E. coli. Nature 290:797–799.

9. Huisman O, D’Ari R, Gottesman S. 1984. Cell-division control in Escherichia coli: specific induction of the SOS function SfiA protein is sufficient to block septation. Proc Natl Acad Sci U S A 81:4490–4494.

10. Jones C, Holland IB. 1985. Role of the SulB (FtsZ) protein in division inhibition during the SOS response in Escherichia coli: FtsZ stabilizes the inhibitor SulA in maxicells. Proc Natl Acad Sci U S A 82:6045–6049.

11. Higashitani A, Higashitani N, Horiuchi K. 1995. A cell division inhibitor SulA of Escherichia coli directly interacts with FtsZ through GTP hydrolysis. Biochem Biophys Res Commun 209:198–204.

12. Mukherjee A, Cao C, Lutkenhaus J. 1998. Inhibition of FtsZ polymerization by SulA, an inhibitor of septation in Escherichia coli. Proc Natl Acad Sci U S A 95:2885–2890.

13. Cordell SC, Robinson E J, Lowe J. 2003. Crystal structure of the SOS cell division inhibitor SulA and in complex with FtsZ. Proc Natl Acad Sci U S A 100:7889–7894.

14. Schoemaker JM, Gayda RC, Markovitz A. 1984. Regulation of cell division in Escherichia coli: SOS induction and cellular location of the sulA protein, a key to Ion-associated filamentation and death. J Bacteriol 158:551–561.

15. Mizusawa S, Gottesman S. 1983. Protein degradation in Escherichia coli: the Ion gene controls the stability of sulA protein. Proc Natl Acad Sci U S A 80:358–362.

16. Modell JW, Hopkins AC, Laub MT. 2011. A DNA damage checkpoint in Caulobacter crescentus inhibits cell division through a direct interaction with FtsW. Genes Dev 25:1328–1343.

17. Modell JW, Kambara TK, Perchuk BS, Laub MT. 2014. A DNA damage-induced, SOS-independent checkpoint regulates cell division in Caulobacter crescentus. PLoS Biol 12:el001977.

18. Tzagoloff H, Novick R. 1977. Geometry of cell division in Staphylococcus aureus. J Bacteriol 129:343–350.

19. Turner RD, Ratcliffe EC, Wheeler R, Golestanian R, Hobbs JK, Foster SJ. 2010. Peptidoglycan architecture can specify division planes in Staphylococcus aureus. Nat Commun 1:26.

20. Pinho MG, Kjos M, Veening JW. 2013. How to get (a)round: mechanisms controlling growth and division of coccoid bacteria. Nat Rev Microbiol 11:601–614.

21. Cirz RT, Jones MB, Gingles NA, Minogue TD, Jarrahi B, Peterson SN, Romesberg FE. 2007. Complete and SOS-mediated response of Staphylococcus aureus to the antibiotic ciprofloxacin. J Bacteriol 189:531–539.

22. Anderson KL, Roberts C, Disz T, Vonstein V, Hwang K, Overbeek R, Olson PD, Projan SJ, Dunman PM. 2006. Characterization of the Staphylococcus aureus heat shock, cold shock, stringent, and SOS responses and their effects on log-phase mRN A turnover. J Bacteriol 188:6739–6756.

23. Cohn MT, Kjelgaard P, Frees D, Penadés JR, Ingmer H. 2011. Clp-dependent proteolysis of the LexA N-terminal domain in Staphylococcus aureus. Microbiology 157:677–684.

24. Mesak LR, Miao V, Davies J. 2008. Effects of subinhibitory concentrations of antibiotics on SOS and DNA repair gene expression in Staphylococcus aureus. Antimicrob Agents Chemother 52:3394–3397.

25. Kawai Y, Moriya S, Ogasawara N. 2003. Identification of a protein, YneA, responsible for cell division suppression during the SOS response in Bacillus subtilis. Mol Microbiol 47: 1113–1122.

26. Buchholz M, Nahrstedt H, Pillukat MH, Deppe V, Meinhardt F. 2013. yneA mRNA instability is involved in temporary inhibition of cell division during the SOS response of Bacillus megaterium. Microbiology 159:1564–1574.

27. van der Veen S, Hain T, Wouters JA, Hossain H, de Vos WM, Abee T, Chakraborty T Wells-Bennik MH. 2007. The heat-shock response of Listeria monocytogenes comprises genes involved in heat shock, cell division, cell wall synthesis, and the SOS response. Microbiology 153:3593–3607.

28. van der Veen S, van Schalkwijk S, Molenaar D, de Vos WM, Abee T, Wells-Bennik MH. 2010. The SOS response of Listeria monocytogenes is involved in stress resistance and mutagenesis. Microbiology 156:374–384.

29. Chauhan A, Lofton H, Maloney E, Moore J, Fol M, Madiraju MV, Rajagopalan M. 2006. Interference of Mycobacterium tuberculosis cell division by Rv2719c, a cell wall hydrolase. Mol Microbiol 62:132–147.

30. Ogino H, Teramoto H, Inui M, Yukawa H. 2008. DivS, a novel SOS-inducible cell-division suppressor in Corynebacterium glutamicum. Mol Microbiol 67:597–608.

31. Buist G, Steen A, Kok J, Kuipers OP. 2008. LysM, a widely distributed protein motif for binding to (peptido)glycans. Mol Microbiol 68:838–847.

32. Mesnage S, Dellarole M, Baxter NJ, Rouget JB, Dimitrov JD, Wang N, Fujimoto Y, Hounslow AM, Lacroix-Desmazes S, Fukase K, Foster SJ, Williamson MP. 2014. Molecular basis for bacterial peptidoglycan recognition by LysM domains. Nat Commun 5:4269.

33. Monteiro JM, Fernandes PB, Vaz F, Pereira AR, Tavares AC, Ferreira MT, Pereira PM, Veiga H, Kuru E, VanNieuwenhze MS, Brun YV, Filipe SR, Pinho MG. 2015. Cell shape dynamics during the staphylococcal cell cycle. Nat Commun 6:8055.

34. Lund VA, Wacnik K, Turner RD, Cotterell BE, Walther CG, Fenn SJ, Grein F, Wollman AJ, Leake MC, Olivier N, Cadby A, Mesnage S, Jones S, Foster SJ. 2018. Molecular coordination of Staphylococcus aureus cell division. Elife 7:e32057.

35. Monteiro JM, Pereira AR, Reichmann NT, Saraiva BM, Fernandes PB, Veiga H, Tavares AC, Santos M, Ferreira MT, Macário V, VanNieuwenhze MS, Filipe SR, Pinho MG. 2018. Peptidoglycan synthesis drives an FtsZ-treadmilling-independent step of cytokinesis. Nature 554:528–532.

36. Vadrevu IS, Lofton H, Sarva K, Blasczyk E, Plocinska R, Chinnaswamy J, Madiraju M, Rajagopalan M. 2011. ChiZ levels modulate cell division process in mycobacteria. Tuberculosis (Edinb) 91:S128–135.

37. Mo AH, Burkholder WF. 2010. YneA, an SOS-induced inhibitor of cell division in Bacillus subtilis, is regulated posttranslationally and requires the transmembrane region for activity. J Bacteriol 192:3159–3173.

38. Tsirigos KD, Peters C, Shu N, Käll L, Elofsson A. 2015. The TOPCONS web server for consensus prediction of membrane protein topology and signal peptides. Nucleic Acids Res 43:W401–7.

39. Karimova G, Ladant D. 2017. Defining Membrane Protein Topology Using pho-lac Reporter Fusions. Methods Mol Biol 1615:129–142.

40. Fey PD, Endres JL, Yajjala VK, Widhelm TJ, Boissy RJ, Bose JL, Bayles KW. 2013. A genetic resource for rapid and comprehensive phenotype screening of nonessential Staphylococcus aureus genes. MBio 4:e00537–12.

41. Carroll RK, Rivera FE, Cavaco CK, Johnson GM, Martin D, Shaw LN. 2014. The lone S41 family C-terminal processing protease in Staphylococcus aureus is localized to the cell wall and contributes to virulence. Microbiology 160:1737–1748.

42. Burby PE, Simmons ZW, Schroeder JW, Simmons LA. 2018. Discovery of a dual protease mechanism that promotes DNA damage checkpoint recovery. PLoS Genet 14:el007512.

43. Steele VR, Bottomley AL, Garcia-Lara J, Kasturiarachchi J, Foster SJ. 2011. Multiple essential roles for EzrA in cell division of Staphylococcus aureus. Mol Microbiol 80:542–555.

44. Bottomley AL, Kabli AF, Hurd AF, Turner RD, Garcia-Lara J, Foster SJ. 2014. Staphylococcus aureus DivlB is a peptidoglycan-binding protein that is required for a morphological checkpoint in cell division. Mol Microbiol 94:1041–1064.

45. Gamba P, Veening JW, Saunders NJ, Hamoen LW, Daniel RA. 2009. Two-step assembly dynamics of the Bacillus subtilis divisome. J Bacteriol 191:4186–4194.

46. Kawai Y, Ogasawara N. 2006. Bacillus subtilis EzrA and FtsL synergistically regulate FtsZ ring dynamics during cell division. Microbiology 152:1129–1141.

47. Pereira SF, Henriques AO, Pinho MG, de Lencastre H, Tomasz A. 2007. Role of PBP1 in cell division of Staphylococcus aureus. J Bacteriol 189:3525–3531.

48. Pereira SF, Henriques AO, Pinho MG, de Lencastre H, Tomasz A. 2009. Evidence for a dual role of PBP1 in the cell division and cell separation of Staphylococcus aureus. Mol Microbiol 72:895–904.

49. den Blaauwen T, Hamoen LW, Levin PA. 2017. The divisome at 25: the road ahead. Curr Opin Microbiol 36:85–94.

50. Daniel RA, Noirot-Gros MF, Noirot P, Errington J. 2006. Multiple interactions between the transmembrane division proteins of Bacillus subtilis and the role of FtsL instability in divisome assembly. J Bacteriol 188:7396–7404.

51. Pinho MG, de Lencastre H, Tomasz A. 2000. Cloning, characterization, and inactivation of the gene pbpC, encoding penicillin-binding protein 3 of Staphylococcus aureus. J Bacteriol 182:1074–1079.

52. Daniel RA, Harry EJ, Katis VL, Wake RG, Errington J. 1998. Characterization of the essential cell division gene ftsL(y 11D) of Bacillus subtilis and its role in the assembly of the division apparatus. Mol Microbiol 29:593–604.

53. Monk IR, Shah IM, Xu M, Tan MW, Foster TJ. 2012. Transforming the untransformable: application of direct transformation to manipulate genetically Staphylococcus aureus and Staphylococcus epidermidis. MBio 3:e00277–11.

54. Bose JL, Fey PD, Bayles KW. 2013. Genetic tools to enhance the study of gene function and regulation in Staphylococcus aureus. Appl Environ Microbiol 79:2218–2224.

55. Monk IR, Tree JJ, Howden BP, Stinear TP, Foster TJ. 2015. Complete Bypass of Restriction Systems for Major Staphylococcus aureus Lineages. MBio 6:e00308-15.

56. Chen J, Yoong P, Ram G, Torres VJ, Novick RP. 2014. Single-copy vectors for integration at the SaPll attachment site for Staphylococcus aureus. Plasmid 76:1–7.

57. Helle L, Kull M, Mayer S, Marincola G, Zelder ME, Goerke C, Wolz C, Bertram R. 2011. Vectors for improved Tet repressor-dependent gradual gene induction or silencing in Staphylococcus aureus. Microbiology 157:3314–3323.

58. Brzoska AJ, Firth N. 2013. Two-plasmid vector system for independently controlled expression of green and red fluorescent fusion proteins in Staphylococcus aureus. Appl Environ Microbiol 79:3133–3136.

59. Wacnik K. 2016. Dissecting cell division in the human pathogen Staphylococcus aureus. PhD Thesis. University of Sheffield.

60. Karimova G, Pidoux J, Ullmann A, Ladant D. 1998. A bacterial two-hybrid system based on a reconstituted signal transduction pathway. Proc Natl Acad Sci U S A 95:5752–5756.

61. Claessen D, Emmins R, Hamoen LW, Daniel RA, Errington J, Edwards DH. 2008. Control of the cell elongation-division cycle by shuttling of PBP1 protein in Bacillus subtilis. Mol Microbiol 68:1029–1046.

62. Bottomley AL, Liew ATF, Kusuma KD, Peterson E, Seidel L, Foster SJ, Harry EJ. 2017. Coordination of Chromosome Segregation and Cell Division in Staphylococcus aureus. Front Microbiol 8:1575.

