## Supplementary data for "SosA inhibits cell division in *Staphylococcus aureus* in response to DNA damage"

**Supplementary Table S1.**

A. Bacterial strains used (*S. aureus* if not otherwise indicated).

| **Name** | **Characteristics** | **Source/**  **Reference** |
| --- | --- | --- |
| *E. coli* DC10B | Lab strain used for cloning | [1] |
| *E. coli* IM08B | Lab strain used for cloning | [2] |
| *E. coli* BTH101 | Δ*cya* strain used for bacterial two-hybrid analysis | [3] |
| RN4220 | Lab strain, resitriction modification-deficient | Lab stock |
| 8325-4 | Lab strain | Lab stock |
| 8325-4Δ*sosA* | Clean deletion of *sosA* in 8325-4 | This study |
| 8325-4Δ*sosA*-compl. | Complementation of 8325-4Δ*sosA* at *attC* site; Erm^R^ | This study |
| 8325-4-*ctpA* | Transposon insertion in *ctpA* of 8325-4 transduced from NE847 [3]; Erm^R^ | This study |
| 8325-4Δ*sosA*-*ctpA* | Transposon insertion in *ctpA* of 8325-4 Δ*sosA* transduced from NE847 [3]; Erm^R^ | This study |
| JE2 | Lab strain | Lab stock |
| JE2Δ*sosA* | Clean deletion of *sosA* in JE2 | This study |
| JE2-*ctpA* | Transposon insertion in *ctpA* of JE2, transduced from NE847 [3]; Erm^R^ | This study |
| JE2-*ctpA*(−erm) | Erm-sensitive derivative of JE2-*ctpA* | This study |
| JE2Δ*sosA*-*ctpA* | Transposon insertion in *ctpA* of JE2 Δ*sosA*, transduced from NE847 [4]; Erm^R^ | This study |
| RN9011 | RN4220/pRN7023 (SaPI1 integrase, cat194) | [5] |
| RN9011-*sosA*-compl. | Integration of pJC1112-*sosA* at *attC* site; Erm^R^ | This study |
| SH4665 | SH1000 pCQ11-FtsZ-eYFP; Erm^R^ | [6] |
| JGL227 | SH1000 *ezrA-gfp*+; Erm^R^ | [7] |
| JGL228 | SH1000 *gpsB -gfp*+; Erm^R^ | [7] |
| SJF4693 | JE2 pRAB12-*lacZ* pCQ11-FtsZ-eYFP; Cm^R^, Erm^R^ | This study |
| SJF4694 | JE2 pSosA pCQ11-FtsZ-eYFP; Cm^R^, Erm^R^ | This study |
| SJF4696 | JE2 pRAB12-*lacZ* *ezrA-gfp*+; Cm^R^, Erm^R^ | This study |
| SJF4697 | JE2 pSosA *ezrA-gfp*+; Cm^R^, Erm^R^ | This study |
| SJF4699 | JE2 pRAB12-*lacZ* *gpsB-gfp*+; Cm^R^, Erm^R^ | This study |
| SJF4700 | JE2 pSosA *gpsB-gfp*+; Cm^R^, Erm^R^ | This study |

B. Plasmids used.

| **Name** | **Characteristics** | **Source/**  **Reference** |
| --- | --- | --- |
| pRAB12-*lacZ* | Expression vector; Cm^R^ | [8] |
| pIMAY | Vector for temperature-sensitive allelic replacement; Cm^R^ | [1] |
| pIMAY-*ΔsosA* | *ΔsosA* deletion fragment cloned into pIMAY; Cm^R^ | This study |
| pSosA | *sosA* cloned behind anhydrotetracycline-inducible promoter of pRAB12-*lacZ;* Cm^R^ | This study |
| pSosAd10 | *sosAd10* cloned behind anhydrotetracycline-inducible promoter of pRAB12-*lacZ;* Cm^R^ | This study |
| pSosAd20 | *sosAd20* cloned behind anhydrotetracycline-inducible promoter of pRAB12-*lacZ;* Cm^R^ | This study |
| pSosAd30 | *sosAd30* cloned behind anhydrotetracycline-inducible promoter of pRAB12-*lacZ;* Cm^R^ | This study |
| pSosAd40 | *sosAd40* cloned behind anhydrotetracycline-inducible promoter of pRAB12-*lacZ;* Cm^R^ | This study |
| pSK9067 | Expression vector; Erm^R^ | [9] |
| pPBP1 | *pbpA* cloned into pSK9067, IPTG-inducible; Erm^R^ | This study |
| pPBP1* | *pbpA** cloned into pSK9067, IPTG-inducible; Erm^R^ | This study |
| pCtpA | *ctpA* cloned into pSK9067, IPTG-inducible; Erm^R^ | This study |
| pDivIC | *divIC* cloned into pSK9067, IPTG-inducible; Erm^R^ | This study |
| pPBP3 | *pbpC* cloned into pSK9067, IPTG-inducible; Erm^R^ | This study |
| pFtsL | *ftsL* cloned into pSK9067, IPTG-inducible; Erm^R^ | This study |
| pMurJ | *murJ* cloned into pSK9067, IPTG-inducible; Erm^R^ | This study |
| pJC1112 | Vector for integration into *attC* site; Erm^R^ | [5] |
| pJC1112-*sosA* | *sosA* behind its own promoter cloned into pJC1112; Erm^R^ | This study |
| pTnT | Vector for elimination of Erm^R^ from transposon insertion | [10] |
| pKT25 | Low copy-number derivative of pSU40 encoding the T25 fragment of *B.pertussis* adenylate cyclase, corresponding to the first 224 amino acids of CyaA, upstream of a multiple cloning site; Kan^R^ | [11] |
| p25-N | Derivative of low-copy number pSU40 encoding the T25 fragment of CyaA, corresponding to the first 224 amino acids, downstream of a multiple cloning site; Kan^R^ | [12] |
| pUT18C | High copy-number derivative of pUC19 encoding the T18 fragment, corresponding to amino acids 225 to 399 of CyaA, upstream of a multiple cloning site; Amp^R^ | [11] |
| pKT25-zip | pKT25 containing the 5’ end of the leucine zipper of GCN4 fused in frame to T25; Kan^R^ | [11] |
| pUT18C-zip | pUT18C containing the 5’ end of the leucine zipper of GCN4 fused in frame to T18; Amp^R^ | [11] |
| pGL540 | pKT25 containing T25 fused in frame to the 5’ end of *S. aureus* *divIB*; Kan^R^ | [7] |
| pGL541 | pKT25 containing T25 fused in frame to the 5’ end of *S. aureus* *ftsA*; Kan^R^ | [7] |
| pGL542 | pKT25 containing T25 fused in frame to the 5’ end of *S. aureus* *ftsL*; Kan^R^ | [7] |
| pGL543 | pKT25 containing T25 fused in frame to the 5’ end of *S. aureus* *pbp2*; Kan^R^ | [7] |
| pGL549 | pKT25 containing T25 fused in frame to the 5’ end of *S. aureus* *ftsZ*; Kan^R^ | [7] |
| pGL550 | pKT25 containing T25 fused in frame to the 5’ end of *S. aureus* *pbpA*; Kan^R^ | [7] |
| pGL551 | pKT25 containing T25 fused in frame to the 5’ end of *S. aureus* *divIC*; Kan^R^ | [7] |
| pGL553 | pKT25 containing T25 fused in frame to the 5’ end of *S. aureus* *parC*; Kan^R^ | [7] |
| pGL554 | pKT25 containing T25 fused in frame to the 5’ end of *S. aureus* *parE*; Kan^R^ | [13] |
| pGL556 | pKT25 containing T25 fused in frame to the 5’ end of *S. aureus* *pbp3*; Kan^R^ | [13] |
| pGL557 | pKT25 containing T25 fused in frame to the 5’ end of *S. aureus* *gpsB*; Kan^R^ | [7] |
| pGL558 | pKT25 containing T25 fused in frame to the 5’ end of *S. aureus* *ypsA*; Kan^R^ | [13] |
| pGL559 | pKT25 containing T25 fused in frame to the 5’ end of *S. aureus* *sepF*; Kan^R^ | [7] |
| pGL560 | pKT25 containing T25 fused in frame to the 5’ end of *S. aureus* *noc*; Kan^R^ | [14] |
| pALB3 | pKT25 containing T25 fused in frame to the 5’ end of *S. aureus* *ftsW*; Kan^R^ | [7] |
| pALB5 | pKT25 containing T25 fused in frame to the 5’ end of *S. aureus* *mreC*; Kan^R^ | [13] |
| pALB8 | pKT25 containing T25 fused in frame to the 5’ end of *S. aureus* *rodA*; Kan^R^ | [7] |
| pALB9 | pKT25 containing T25 fused in frame to the 5’ end of *S. aureus* *zapA*; Kan^R^ | [13] |
| pVF30 | p25-N containing T25 fused in frame to the 3’ end *of S. aureus ezrA*; Kan^R^ | [7] |
| pALB50 | p25-N containing T25 fused in frame to the 3' end of *S. aureus divIVA*; Kan^R^ | [15] |
| pT18-SosA | pUT18C containing T18 fused in frame to the 5’ end of *S. aureus* *sosA*; Amp^R^ | This study |
| pT18-SosA(44A) | pUT18C containing T18 fused in frame to the 5’ end of *S. aureus* *sosA(44A)*; Amp^R^ | This study |
| pT18-SosAd10 | pUT18C containing T18 fused in frame to the 5’ end of *S. aureus* *sosAd10*; Amp^R^ | This study |
| pT18-SosAd40 | pUT18C containing T18 fused in frame to the 5’ end of *S. aureus* *sosAd40*; Amp^R^ | This study |
| pSosAd10(37A) | Alanine substitution variant of SosAd10 cloned into pRAB12-*lacZ*; Cm^R^ | This study |
| pSosAd10(37A/38A) | Alanine substitution variant of SosAd10 cloned into pRAB12-*lacZ*; Cm^R^ | This study |
| pSosAd10(38A) | Alanine substitution variant of SosAd10 cloned into pRAB12-*lacZ*; Cm^R^ | This study |
| pSosAd10(40A) | Alanine substitution variant of SosAd10 cloned into pRAB12-*lacZ*; Cm^R^ | This study |
| pSosAd10(40A/41A) | Alanine substitution variant of SosAd10 cloned into pRAB12-*lacZ*; Cm^R^ | This study |
| pSosAd10(41A) | Alanine substitution variant of SosAd10 cloned into pRAB12-*lacZ*; Cm^R^ | This study |
| pSosAd10(44A) | Alanine substitution variant of SosAd10 cloned into pRAB12-*lacZ*; Cm^R^ | This study |
| pSosAd10(44A/45A) | Alanine substitution variant of SosAd10 cloned into pRAB12-*lacZ*; Cm^R^ | This study |
| pSosAd10(45A) | Alanine substitution variant of SosAd10 cloned into pRAB12-*lacZ*; Cm^R^ | This study |
| pSosA(44A) | Alanine substitution variant of full length SosA in pRAB12-*lacZ*; Cm^R^ | This study |
| pKTop | Vector with *phoA-lacZ* fusion for membrane topology analysis; Kan^R^ | [16] |
| pKTop-s*osA* | *sosA* cloned in frame into pKTop; Kan^R^ | This study |
| pKTop-*sosAd10* | *sosAd10* cloned in frame into pKTop; Kan^R^ | This study |
| pKTop-*sosAd10(44A)* | *sosAd10(44A)* cloned in frame into pKTop; Kan^R^ | This study |
| pKTop-*sosAd40* | *sosAd40* cloned in frame into pKTop; Kan^R^ | This study |

C. Oligonucleotides used.

| **Name** | **Sequence (5’-3’)** |
| --- | --- |
| Up-sosA_fw-KpnI | ATATGGTACCCTCGCTCCTGTAAATTATTACG |
| Up-sosA_rev | TTTCACTCCTAGAACATTTGTTTG |
| Dw-sosA_fw | CAAATGTTCTAGGAGTGAAATACATTGTCACAACGTTATTTTG |
| Dw-sosA_rev-SacI | ATATGAGCTCCATATGTGTAATGATCTACAACATTATATC |
| Ctrl_dsosA_F | ATTCTCTCATATATAGGCACTCC |
| Ctrl_dsosA_R | CTGTTTGCTCCTTTGCTTC |
| Fwd_MCS | TACATGTCAAGAATAAACTGCCAAAGC |
| Rev_MCS | AATACCTGTGACGGAAGATCACTTCG |
| ctpA_F-SalI | ATATGTCGACCATAATAAGGAAGTGATACAATGG |
| ctpA_R-EcoRI | GATACAGAATTCTACAATTTTAGTAGTGTGTATCGC |
| pbpA_F-SalI | ATATGTCGACGAGAACGATAATGTAAAGGTAGTG |
| pbpA_R-EcoRI | GATACAGAATTCTTAGTCCGACTTATCCTTGTC |
| divIC-F_SalI | ATATGTCGACTAAATTGGAGGTGACAAGCAATG |
| divIC-R_EcoRI | GATACAGAATTCTTATTTTTTCGAAGATTTTGAGCTAGAC |
| pbpC-F_SalI | ATATGTCGACCTTTGAATAGAGGTAGGTAGTTTTG |
| pbpC-R_EcoRI | GATACAGAATTCTTATTTGTCTTTGTCTTTATTTTTATCATC |
| ftsL-F_SalI | ATATGTCGACTACTTAAATAAGGAGCAATTTATAATGG |
| ftsL-R_AatII | GATACAGACGTCTTAATTTTTTGCTTCGCCATTACTAC |
| murJ-F_SalI | ATATGTCGACATGAGATAGGGAGATTCGTAATG |
| murJ-R_AatII | GATACAGACGTCTCATCGTAAAAACCTAACTCTAC |
| Up-sosA-promo_SalI | GATACAGTCGACCTCTCATATATAGGCACTCCC |
| Up-sosA_BglII | GATACAAGATCTGTTCTAGGAGTGAAAATGATG |
| Dw-sosA_EcoRI | GATACAGAATTCTCAATTTATTAAAGCGAACAC |
| Dw-sosA(d10)_EcoRI | GATACAGAATTCTCATTGTTCGCTATTGTTTGTAG |
| Dw-sosA(d20)_EcoRI | GATACAGAATTCTCATTCGTATGCTTTATTTATCGT |
| Dw-sosA(d30)_EcoRI | GATACAGAATTCTCAAATTTGATGGTCAGTCATTTC |
| Dw-sosA(d40)_EcoRI | GATACAGAATTCTCATTCCGAGTGAGCACTAATG |
| SosA_R-long | ATTTATTAAAGCGAACACTTTCCCATCTCTTTGTTCGCTATTGTTTGTAG |
| SosA_F-BamHI | GATACAGGATCCCATGTTTTACAATAAATATAAAAACG |
| SosA_R-KpnI | GATACAGGTACCTCATTTATTAAAGCGAACACTTT |
| SosAd10_R-KpnI | GATACAGGTACCTCTTGTTCGCTATTGTTTGTAG |
| SosAd40_R-KpnI | GATACAGGTACCTCTTCCGAGTGAGCACTAATG |
| ALB133 | AATTAAGGATCCAATGTTTTACAATAAATATAAAAACGT |
| ALB134 | AAAAATGAATTCTCAATTTATTAAAGCGAACACT |

**Supplementary Figure S1.**

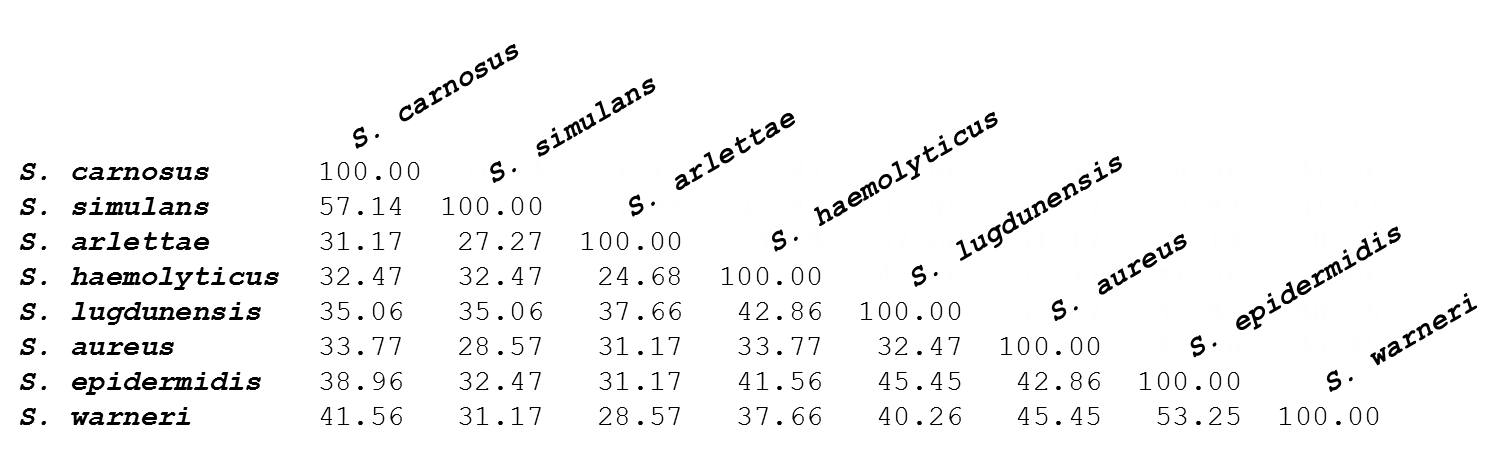

**Fig. S1** Percent identity matrix for staphylococcal SosA proteins. Pairwise identity scores for the proteins included in the alignment in Figure 1 were obtained by the Clustal 2.1 algorithm.

**Supplementary Figure S2.**

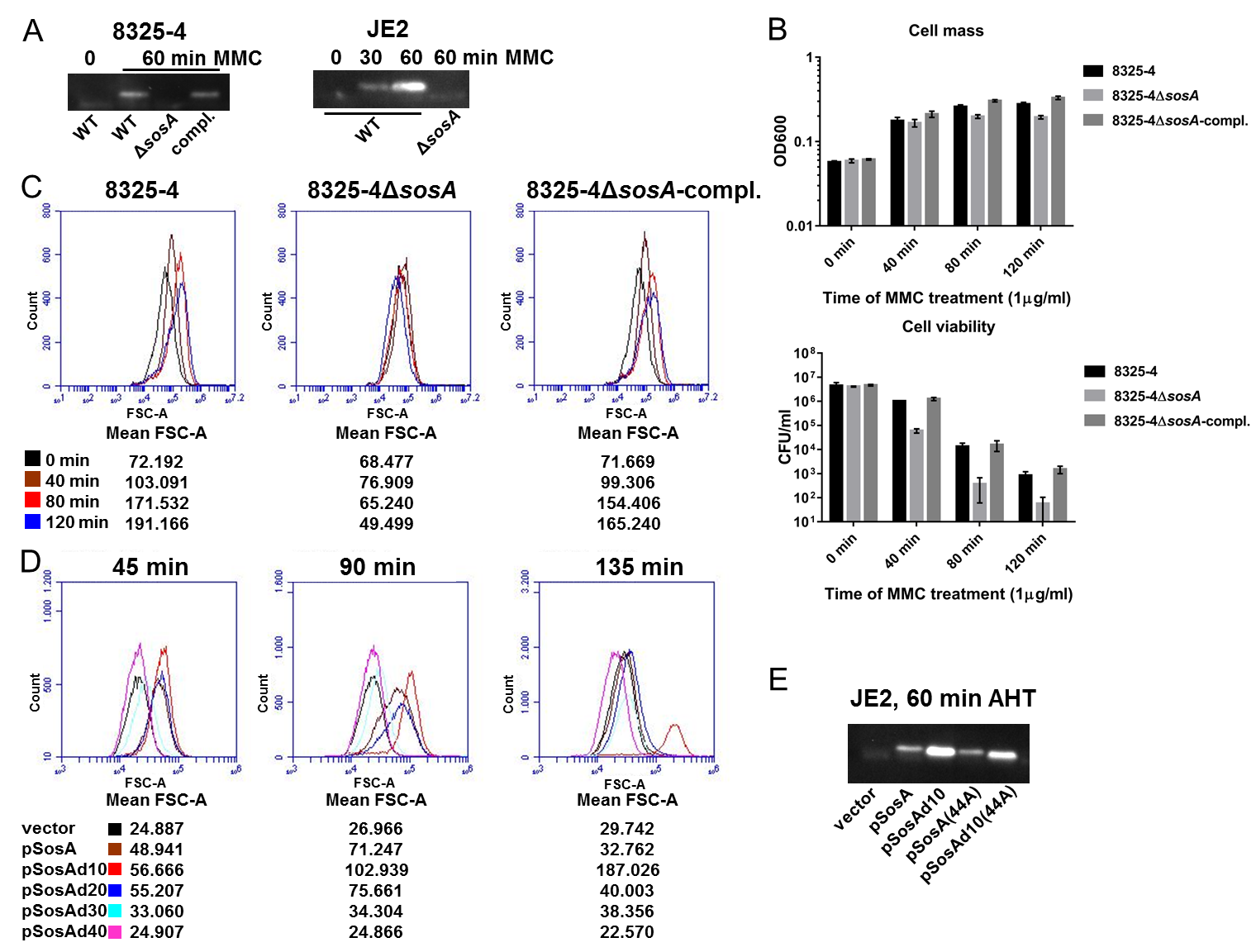

**Fig. S2** (A) Detection by western blot of DNA-damage induced expression of SosA. SosA being detected in wildtype *S. aureus* 8325-4 and the complemented strain, while being absent in the *sosA* deletion mutant following 1 h of MMC treatment (1 µg/ml). Also displayed is the time-dependent expression of SosA in *S. aureus* JE2 post MMC addition (1 µg/ml) in comparison to the corresponding *sosA* deletion mutant. (B and C) Phenotypic complementation of the *sosA* deletion mutant. The phenotypes of *S. aureus* 8325-4Δ*sosA* were restored back to wildtype by chromosomal integration of a copy of the *sosA* gene under its native promoter (*S. aureus* 8325-4Δ*sosA*-compl.) when evaluated for changes in cell density and viability (B) and cell size assessed by flow cytometry (C) during challenge with MMC (1 µg/ml). Mean FSC-A values are indicated below histograms. Cells were grown exponentially prior to addition of MMC at an OD_600_ of 0.05. (D) Effect of different truncated SosA variants on *S. aureus* cell size. Evaluation of cell size distribution was performed by flow cytometry (FSC-A) of *S. aureus* RN4220 containing expression plasmids encoding full length SosA or C-terminally truncated variants of the protein. Mean FSC-A values are indicated below histograms. Cells were grown exponentially prior to induction with 100 ng/ml of AHT and analyzed at indicated time points. (E) Western blot of accumulation in *S. aureus* JE2 of SosA, SosAd10, SosA(44A), and SosAd10(44A) expressed from respective plasmid constructs for 1 h with 200 ng/ml AHT.

**Supplementary Figure S3.**

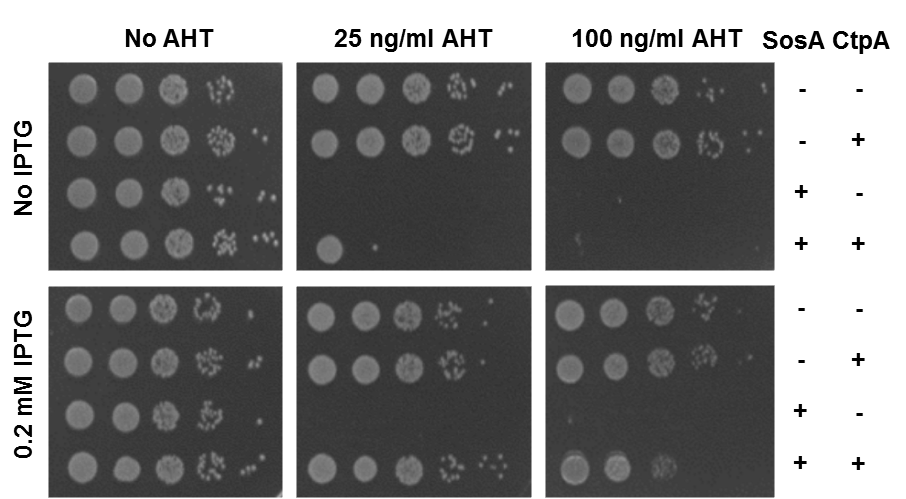

**Fig. S3** Hypersusceptibility of an *S. aureus* JE2 *ctpA* mutant to SosA-mediated growth inhibition. Plating efficiency of the *S. aureus* JE2-*ctpA*(−erm) mutant transformed with both SosA and CtpA expression plasmids (+) or vector controls (-) at different inducer concentrations; AHT for SosA and IPTG for CtpA. Cells were grown exponentially to an OD_600_ of 0.5, serially 10-fold diluted, and plated on TSA plus indicated inducer concentrations followed by incubation overnight at 37°C before imaging.

**Supplementary Figure S4.**
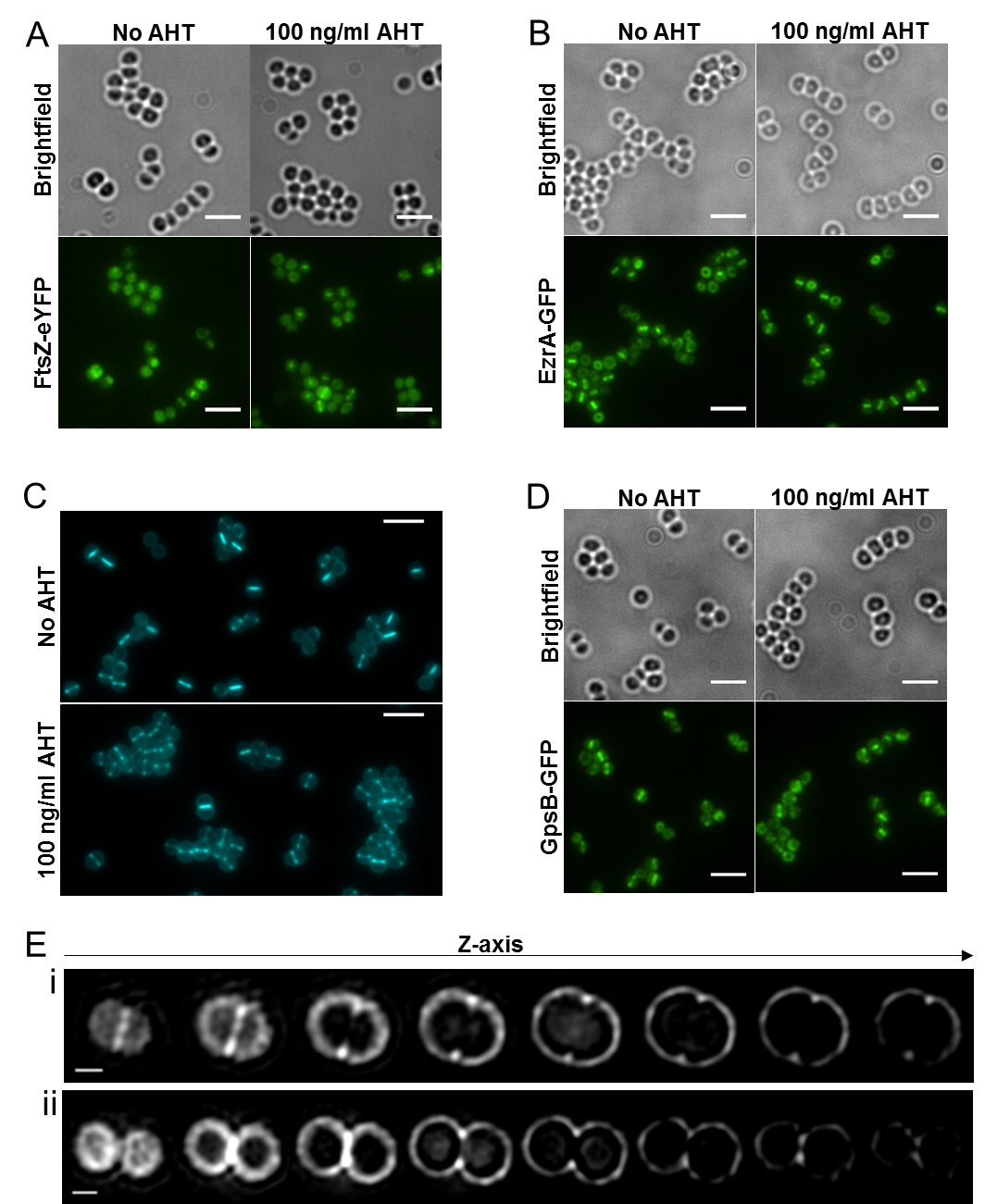

**Fig. S4** SosA halts septum completion. Localization of (A) FtsZ-eYFP in SJF4693 (JE2 pRAB12-*lacZ* pCQ11-FtsZ-eYFP ), (B) EzrA-GFP in SJF4696 (JE2 pRAB12-*lacZ* *ezrA-gfp*+) and (D) GpsB-GFP in SJF4699 (JE2 pRAB12-*lacZ* *gpsB-gfp*+) grown in the absence and presence of 100 ng/ml of AHT for 45 min. Fluorescence images are average intensity projections. Scale bars represents 3 µm. (C) Fluorescence microscopy images of JE2/pSosA grown in the absence or presence of 100 ng/ml of AHT for 45 min and labeled with HADA for 5 min. Images are average intensity projections. Scale bars represents 3 µm. (E) 3D-SIM Z-stack images of JE2/pSosA grown with 100 ng/ml of AHT for 45 min and labelled with Alexa Fluor 647 NHS ester. (i) A cell with initiated septum formation and (ii) a cell splitting into daughter cells without finishing a septal disc. Scale bars represents 0.5 µm.

**Supplementary Figure S5.**

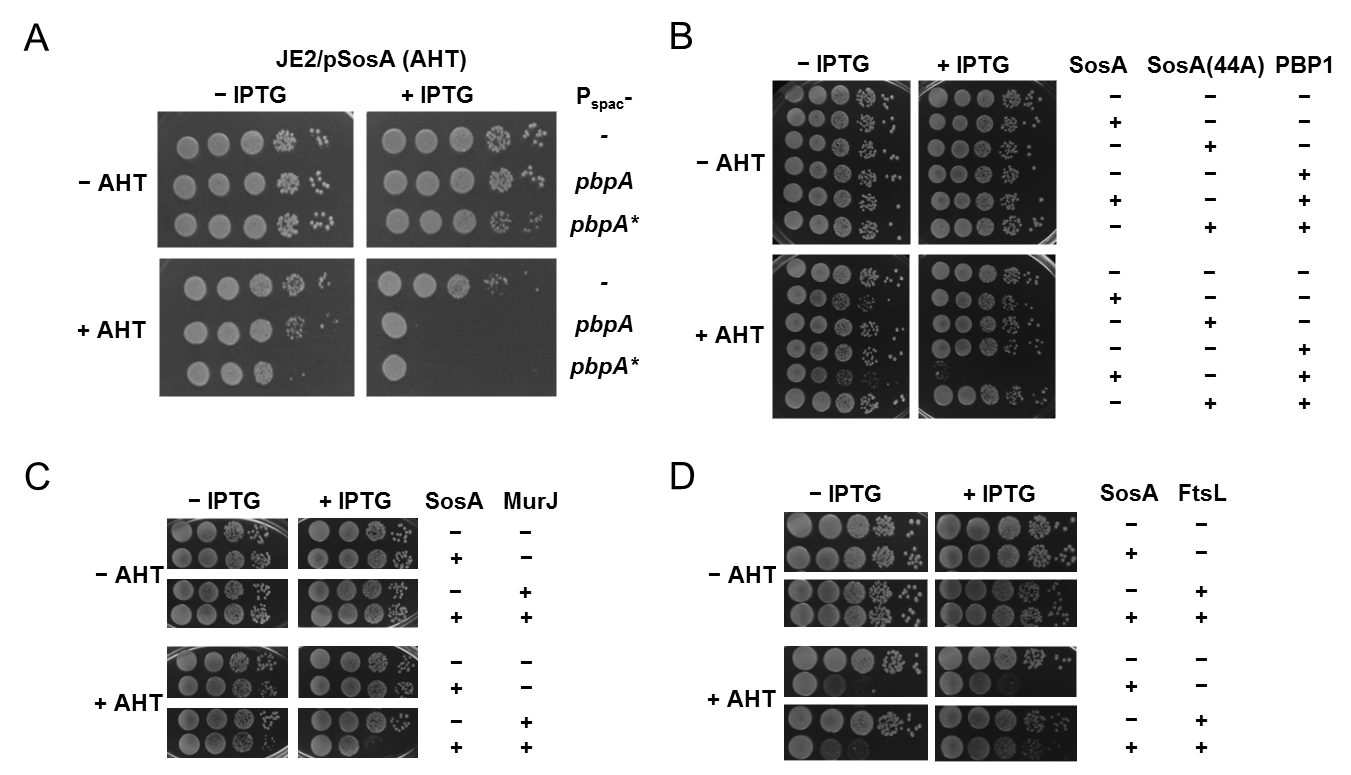

**Fig. S5** Assessment of the plating efficiency of *S. aureus* strain JE2 upon expression of *sosA* (pSosA, AHT-inducible) and concomitant overexpression of PBP1 (*pbpA*) or a PBP1 transpeptidase mutant PBP1* (*pbpA**) (A), PBP1 (*pbpA*) and in comparison to SosA(44A) (B), MurJ (C), and FtsL (D). Inducer concentrations for SosA (and SosA(44A)) were 50 ng/ml (panels A-C) and 150 ng/ml AHT (panel D). Co-expression proteins were induced by 0.2 mM IPTG. (+) denotes protein encoding vectors, (-) denotes empty vector controls. In all panels, cells were grown exponentially to an OD_600_ of 0.5, serially 10-fold diluted, and plated on TSA with/without inducers followed by incubation overnight at 37°C before imaging.

**Supplementary Figure S6.**

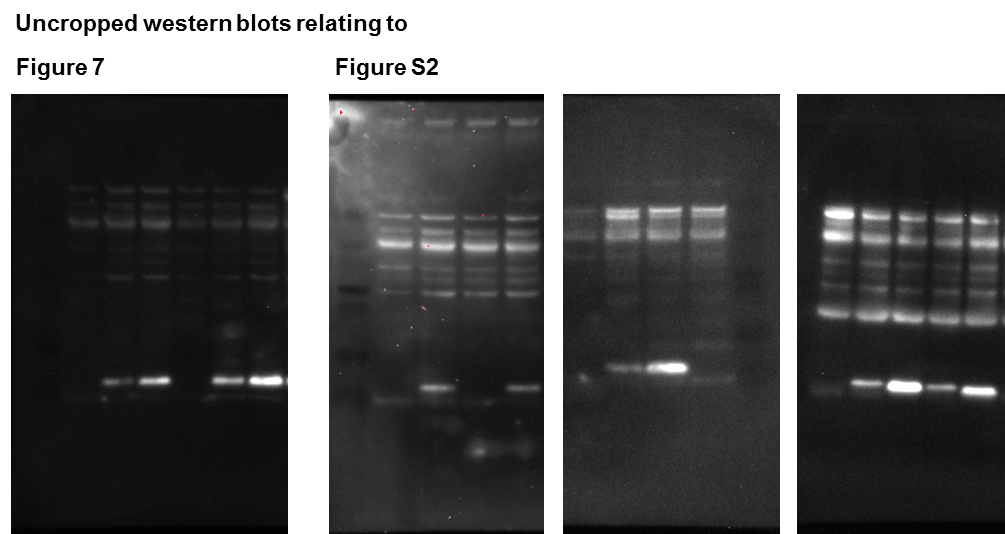

**Fig. S6** Uncropped western blots relating to Figure 7 and Figure S2.

**Supplementary Movie S1.**

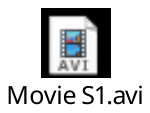
Time-lapse microscopy of *S. aureus* JE2 WT and *sosA* mutant upon MMC exposure.

**Supplementary Movie S2.**

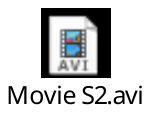
Time-lapse microscopy of *S. aureus* JE2 cells overexpressing SosA.

**Supplementary Movie S3.**

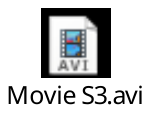
Time-lapse microscopy of *S. aureus* JE2 cells overexpressing SosAd10.

**References for Supplemental Material**

1. Monk IR, Shah IM, Xu M, Tan MW, Foster TJ. 2012. Transforming the untransformable: application of direct transformation to manipulate genetically Staphylococcus aureus and Staphylococcus epidermidis. MBio 3:e00277-11.
2. Monk IR, Tree JJ, Howden BP, Stinear TP, Foster TJ. 2015. Complete Bypass of Restriction Systems for Major Staphylococcus aureus Lineages. MBio 6:e00308-15.
3. Karimova G, Ladant D. 2005. A bacterial two-hybrid system based on a Cyclic AMP signaling cascade. Chap. 26, pp 499-515. Protein-Protein Interactions, A Molecular Cloning Manual 2nd Edition. Cold Spring Harbor Laboratory Press. Edited by E. Golemis. Cold Spring Harbor, New York.
4. Fey PD, Endres JL, Yajjala VK, Widhelm TJ, Boissy RJ, Bose JL, Bayles KW. 2013. A genetic resource for rapid and comprehensive phenotype screening of nonessential Staphylococcus aureus genes. MBio 4:e00537-12.
5. Chen J, Yoong P, Ram G, Torres VJ, Novick RP. 2014. Single-copy vectors for integration at the SaPI1 attachment site for Staphylococcus aureus. Plasmid 76:1-7.
6. Lund VA, Wacnik K, Turner RD, Cotterell BE, Walther CG, Fenn SJ, Grein F, Wollman AJ, Leake MC, Olivier N, Cadby A, Mesnage S, Jones S, Foster SJ. 2018. Molecular coordination of Staphylococcus aureus cell division. Elife 7:e32057.
7. Steele VR, Bottomley AL, Garcia-Lara J, Kasturiarachchi J, Foster SJ. 2011. Multiple essential roles for EzrA in cell division of Staphylococcus aureus. Mol Microbiol 80:542-555.
8. Helle L, Kull M, Mayer S, Marincola G, Zelder ME, Goerke C, Wolz C, Bertram R. 2011. Vectors for improved Tet repressor-dependent gradual gene induction or silencing in Staphylococcus aureus. Microbiology 157:3314-3323.
9. Brzoska AJ, Firth N. 2013. Two-plasmid vector system for independently controlled expression of green and red fluorescent fusion proteins in Staphylococcus aureus. Appl Environ Microbiol 79:3133-3136.
10. Bose JL, Fey PD, Bayles KW. 2013. Genetic tools to enhance the study of gene function and regulation in Staphylococcus aureus. Appl Environ Microbiol 79:2218-2224.
11. Karimova G, Pidoux J, Ullmann A, Ladant D. 1998. A bacterial two-hybrid system based on a reconstituted signal transduction pathway. Proc Natl Acad Sci U S A 95:5752-5756.
12. Claessen D, Emmins R, Hamoen LW, Daniel RA, Errington J, Edwards DH. 2008. Control of the cell elongation-division cycle by shuttling of PBP1 protein in Bacillus subtilis. Mol Microbiol 68:1029-1046.
13. Bottomley AL. 2011. Identification and characterisation of the cell division machinery in Staphylococcus aureus. University of Sheffield.
14. Fairclough V. 2009. Functional analysis of novel essential genes of *Staphylococcus aureus*. PhD Thesis. University of Sheffield.
15. Bottomley AL, Liew ATF, Kusuma KD, Peterson E, Seidel L, Foster SJ, Harry EJ. 2017. Coordination of Chromosome Segregation and Cell Division in Staphylococcus aureus. Front Microbiol 8:1575.
16. Karimova G, Ladant D. 2017. Defining Membrane Protein Topology Using pho-lac Reporter Fusions. Methods Mol Biol 1615:129-142.
